# Circular RNA circASH1L(4,5) protects microRNA-129-5p from target-directed microRNA degradation in human skin wound healing

**DOI:** 10.1101/2024.03.12.584463

**Authors:** Qizhang Wang, Guanglin Niu, Zhuang Liu, Maria A. Toma, Jennifer Geara, Xiaowei Bian, Letian Zhang, Minna Piipponen, Dongqing Li, Pehr Sommar, Ning Xu Landén

## Abstract

Both circular RNAs (circRNA) and microRNAs (miRNA) have emerged to play important roles in health and disease. To understand their function in tissue repair, we profiled circRNA, linear RNA, and miRNA expression dynamics in human wound-edge keratinocytes across the wound healing process. Our investigation spotlighted circASH1L(4,5) and its engagement with miR-129-5p, both of which levels were increased with wound repair. Unlike conventional miRNA sponging, circASH1L enhanced miR-129 stability and silencing activity by protecting this miRNA from target-directed miRNA degradation (TDMD) triggered by NR6A1 mRNA. TGF-β signaling, pivotal in wound healing, fostered circASH1L expression while suppressing NR6A1, thus enhancing the miR-129 abundance at the post-transcriptional level. Functionally, circASH1L and miR-129 enhanced keratinocyte migration and proliferation, crucial for re-epithelialization of human wounds. Collectively, our study uncovers circRNAs’ novel role as shields for miRNAs and sheds light on the physiological importance of regulated miRNA degradation in human skin wound healing.

## Introduction

After skin injury, keratinocytes at the wound edge promptly undergo migration and proliferation for re-epithelialization ^1^. These fundamental biological events are underpinned by a well-orchestrated gene expression program that remains largely undisclosed in humans. Failed re-epithelialization is a common pathological hallmark in all types of chronic ulcers, which have posed a major and increasing health and economic burden worldwide ^1,2^. An in-depth comprehension of the gene expression regulation in human skin wound repair is essential for advancing more targeted and effective wound treatments.

Circular RNA (circRNA) represents a novel RNA class with covalently linked 3′ and 5′ ends, exhibiting higher stability due to its loop structure ^3^. They demonstrate diverse functional roles, such as transcriptional regulation, protein scaffolds, translation into proteins, and interaction with microRNAs (miRNAs) ^3^. Among these, miRNA sponging is the most reported model — circRNAs, functioning as competing endogenous RNAs (ceRNAs), bind to miRNAs, preventing the suppression of miRNA targets ^3^. However, the fate of miRNAs post-binding to circRNAs remains largely unexplored.

Recent discoveries have unveiled target-directed miRNA degradation (TDMD) as a prominent mechanism actively and selectively inducing miRNA degradation in animals ^4^. In TDMD, targets that extensively base-pair with both the 5′ and 3′ ends of the miRNA and possess central mismatches trigger a conformational change in Argonaute (AGO) proteins. This change leads to AGO’s poly-ubiquitination by ZSWIM8 E3 ubiquitin ligase and subsequent degradation by the proteasome, rendering the loaded miRNAs unprotected and susceptible to degradation by RNases ^5,6^. Notably, circRNA Cdr1as, the first circRNA that has been studied functionally, can shield miRNA-7 from Cyrano (a long noncoding RNA)-mediated TDMD, which constitutes of a network of noncoding RNAs important for mouse brain functions ^7-9^. Lately, Wightman F.F. *et. al.* proposed a model that circRNAs, unlike linear mRNAs, led to a topology-dependent TDMD evasion, aiding in the stabilization of specific miRNAs^10^. These findings position circRNAs as novel contributors to the endogenous TDMD mechanism. However, the physiological relevance of circRNA-mediated miRNA stabilization remains to be determined.

Both circRNAs and miRNAs have emerged as significant contributors to skin wound healing^11^. To gain insight into their expression and potential interplays during human skin wound healing, we have profiled the *in vivo* expression changes of circRNA, mRNA, and miRNA across the healing process in humans ^12,13^. Our comprehensive dataset revealed that circASH1L(4,5) (circBase: hsa_circ_0003247) ^14,15^ carries four binding sites for miR-129-5p, and the expression of these two RNAs increased in epidermal cells during wound repair. In contrast to the commonly reported miRNA sponging model, circASH1L(4,5) enhanced miR-129-5p stability and function by safeguarding it from NR6A1 mRNA-triggered TDMD. We provided compelling evidence demonstrating the functional significance of this novel circASH1L/miR-129-5p axis in promoting human keratinocyte migration, proliferation, and wound re-epithelialization.

## Results

### CircASH1L(4,5) is up-regulated in keratinocytes during human skin wound healing

To examine the *in vivo* gene expression dynamics during human skin wound healing, we created wounds on the skin of 15 healthy volunteers and collected skin and wound-edge tissues on post-wounding day one (W1) and day seven (W7) from the same donors, representing the inflammatory and proliferative phases of wound healing, respectively ^16^. Paired small RNA sequencing (seq) and ribosomal RNA-depleted long RNA-seq were performed to analyze miRNA, mRNA, and circRNA expression in human skin, W1, and W7 biopsies (n=5 donors) and in epidermal keratinocytes isolated from skin and W7 tissues (n=5 donors, **Fig. 1A**) ^12,13^.

**Figure 1.**
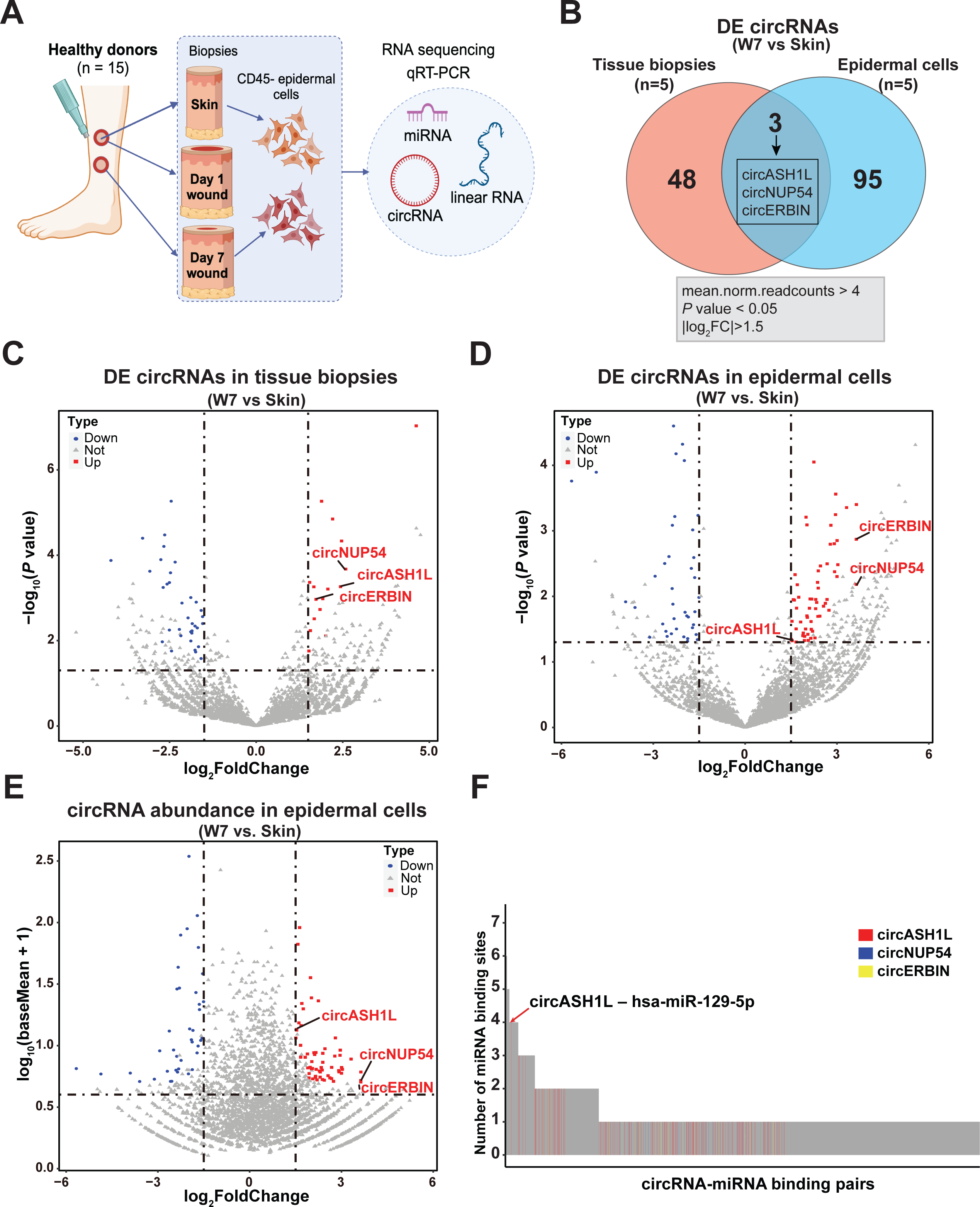
Analysis of circRNA expression changes during human skin wound healing. (A) RNA sequencing and qRT-PCR were conducted in tissue samples and isolated epidermal cells from matched skin, day 1, and day 7 acute wounds of 15 healthy donors. (B) Venn diagram illustrating circRNAs that are differentially expressed (DE) in both day-7 wound (W7) tissues and epidermal cells, relative to the corresponding skin tissues and cells, as shown by RNA-seq. Volcano plots showing the DE circRNAs in W7 tissues compared to the skin (n=5 donors) (C) and in W7 epidermal cells compared to skin cells (n=5 donors) (D). The up- and down-regulated genes are highlighted in red and blue, respectively. (E) Scatter plot showing the abundance and log_2_FC of the DE circRNAs in W7 epidermal cells compared to skin cells. (F) miRNA binding sites were predicted in the 145 DE circRNAs (Fig. 1B). Bar chart showing number of miRNA binding sites for each miRNA-circRNA pair.

Comparing day-7 wounds with donor-matched skin, we identified 51 and 98 differentially expressed [DE: mean normalized read counts of back-spliced junction (BSJ) of circRNAs >4, |log_2_ fold change (FC)| > 1.5, P-value < 0.05] circRNAs in tissue biopsies and isolated epidermal cells, respectively (**Fig. 1B and Table S1**). Three DE circRNAs were identified in both tissue and epidermal cell RNA-seq analyses (**Fig. 1C and 1D**). Among them, circASH1L(4,5) (circBase: hsa_circ_0003247) was notable for its higher expression in epidermal cells compared to the other two DE circRNAs (**Fig. 1E**). Further analysis using Targetscan ^17^ and RNA22 ^18^ to identify miRNA target sites within these DE circRNAs revealed that circASH1L contains four binding sites for hsa-miR-129-5p (**Fig. 1F and 3A**). The combination of its high expression level and the presence of multiple binding sites for the same miRNA underscores circASH1L’s potential as a miRNA sponge.

**Figure 2.**
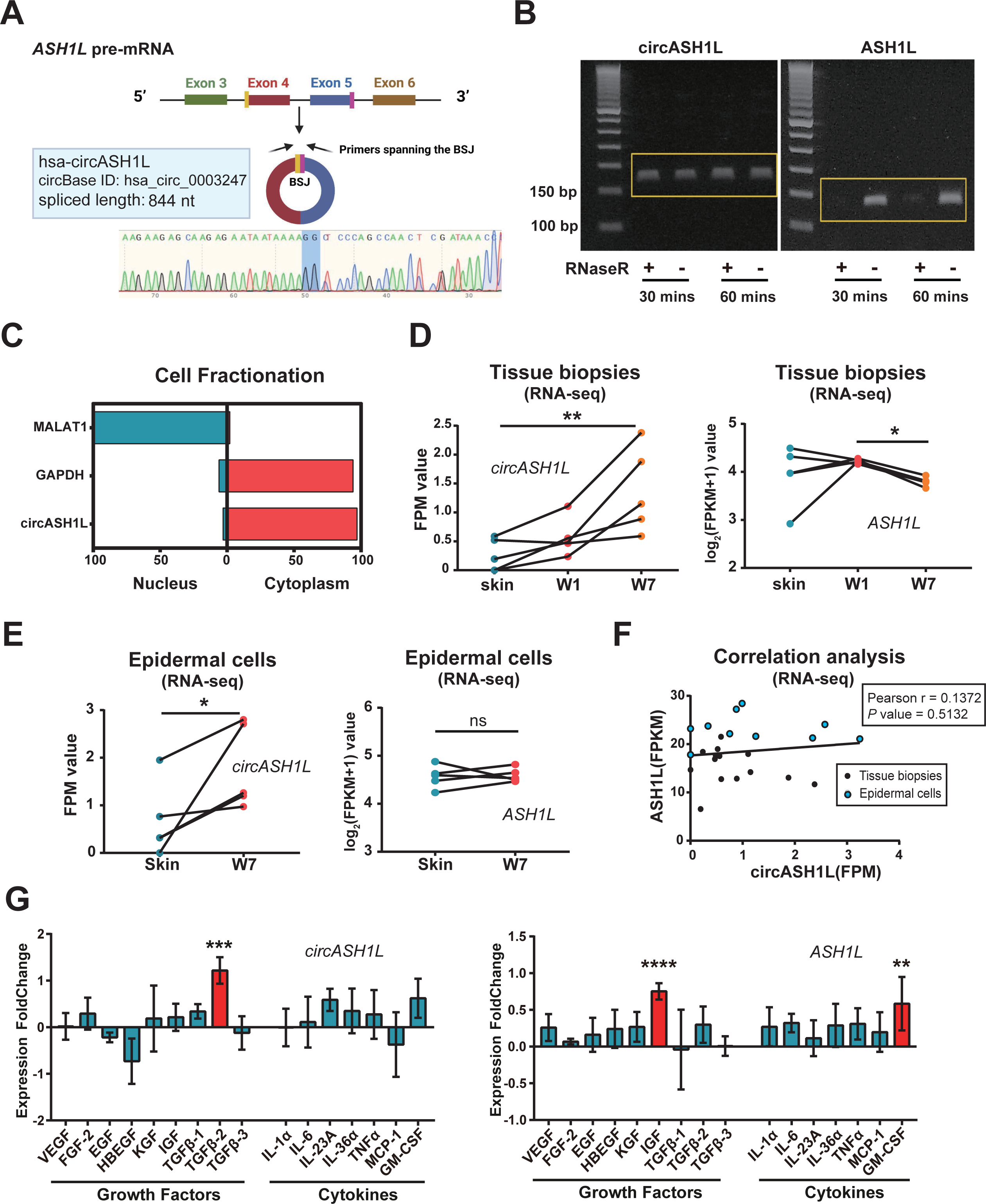
CircASH1L(4,5) is up-regulated in keratinocytes during human skin wound healing. (A) Illustration of circASH1L(4,5) biogenesis. Sanger sequencing verified the back-splicing junction (BSJ) sequence of circASH1L. (B) RT-PCR analysis of circASH1L and linear ASH1L transcripts in human primary keratinocyte (HEKa) RNA treated with or not with RNase R for 30 or 60 minutes. (C) qRT-PCR of circASH1L, GAPDH, and MALAT1 in nuclear and cytoplasmic fractions of HEKa. (D and E) RNA-seq analysis of circASH1L (left) and linear ASH1L (right) in human skin, day 1 (W1), and day 7 wound (W7) tissues (n = 5 donors) (D) and epidermal cells (n=5 donors) (E). (F) Correlation between linear ASH1L and circASH1L expression in human wound tissues and epidermal cells analyzed by RNA-seq (n=25). (G) qRT-PCR analysis of circASH1L (left) and linear ASH1L (right) expression in HEKa treated with wound healing related cytokines and growth factors for 24 hours (n=4). Data are presented as means ± SD or individual values; \**P* < 0.05; \*\**P* < 0.01; \*\*\**P* < 0.001; \*\*\*\**P* < 0.0001; ns, not significant; Paired Student’s t-test (E); One-way ANOVA and multiple comparisons test (D, G); Pearson’s correlation test (F).

**Figure 3.**
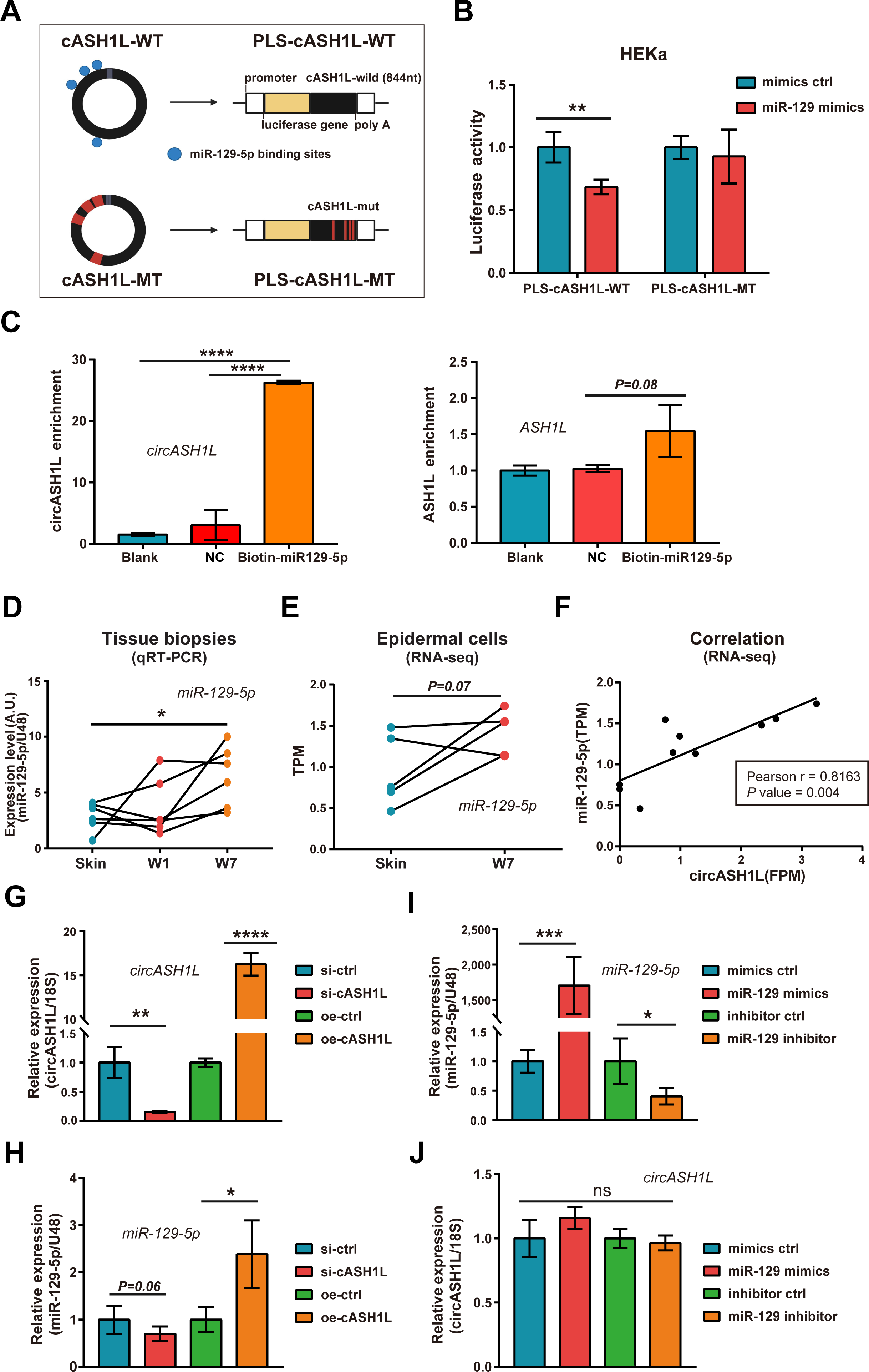
CircASH1L binds to miR-129-5p to increase its level in human wound-edge keratinocytes. (A) Illustration of luciferase reporter plasmids containing wild type (WT) or mutant type (MT) of circASH1L sequence. (B) Luciferase assay was conducted using HEKa cells transfected with the reporter plasmids together with miR-129 mimics or control oligos (n=4). (C) RNA pull-down assay was performed in HEKa transfected with biotin-labelled miR-129-5p, or biotin-labelled control oligos (NC), or without transfection (Blank). CircASH1L (left) and linear ASH1L (right) were detected in the pulled-down RNA by qRT-PCR (n=3). (D) qRT-PCR of miR-129-5p expression in human skin, day 1 (W1), and day 7 wound (W7) tissues (n=7). (E) Paired scatter diagram showing miR-129-5p expression in skin and W7 epidermal cells as shown in small RNA sequencing (n=5 donors). (F) Correlation between miR-129-5p and circASH1L expression in human skin and wound epidermal cells analyzed by RNA-seq (n=10). qRT-PCR analysis of circASH1L (G, J) and miR-129-5p (H, I) in HEKa transfected with circASH1L siRNAs (si-cASH1L) or overexpressing plasmids (oe-cASH1L) or their controls (si-ctrl or oe-ctrl) (n=4) (G, H); and in HEKa transfected with miR-129 mimics, inhibitors, or their controls (n=4) (I, J). Data are presented as means ± SD or individual values; \**P* < 0.05; \*\**P* < 0.01; \*\*\**P* < 0.001; \*\*\*\**P* < 0.0001; ns, not significant. Unpaired Student’s t-test (B, G-J); Paired Student’s t-test (E); One-way ANOVA and multiple comparisons test (C, D); Pearson’s correlation test (F).

CircASH1L(4,5) originates from the 4^th^ and 5^th^ exons of the *ASH1 like histone lysine methyltransferase* (*ASH1L*) gene (**Fig. 2A**). To confirm the circular nature of circASH1L(4,5), we conducted reverse transcription polymerase chain reaction (RT-PCR) using outward-facing primers on total RNA extracted from human primary keratinocytes. Subsequent sequencing of the RT-PCR products validated the sequence of the back-splicing junction (BSJ) of circASH1L(4,5) (**Fig. 2A**). To further substantiate the circularity of circASH1L(4,5), we subjected RNA samples to treatment with RNase R. This enzymatic treatment efficiently digested linear ASHIL transcripts while leaving circASH1L(4,5) unaffected (**Fig. 2B**) ^19^. Additionally, cellular fractionation assays demonstrated that circASH1L, similar to the majority of exonic circRNAs, is predominantly localized in the cytoplasm (**Fig. 2C**) ^20^.

Through RNA-seq analysis, we observed a gradual increase in circASH1L levels during human skin wound healing (**Fig. 2D**), which result was further confirmed by using qRT-PCR in additional human samples (**Fig. S1**). Notably, the expression of circASH1L was significantly elevated in W7 keratinocytes compared to skin keratinocytes from the same donors (**Fig. 2E**). In contrast, the expression of linear ASH1L RNA did not show a similar upregulation during wound healing, and its level at day 7 (W7) was even lower than at day 1 (W1) (**Fig. 2D and 2E**). Also, there was no significant correlation between circASH1L and linear ASH1L expression in these human skin and wound samples, suggesting that circASH1L expression is independent of its cognate linear RNA and may be subject to specific regulatory mechanisms (**Fig. 2F**). To explore this further, we treated human primary keratinocytes with a panel of cytokines and growth factors known to be crucial for wound repair. Our findings revealed that circASH1L expression was significantly enhanced by TGF-β2, while ASH1L mRNA levels were increased by IGF and GM-CSF in keratinocytes (**Fig. 2G**). Despite previous studies demonstrating the requirement of ASH1L for epidermal homeostasis and wound repair ^21^, our data suggest that circASH1L may fulfill functional roles independent of linear ASH1L transcripts during human skin wound healing.

### CircASH1L binds to miR-129-5p to increase its level in human wound-edge keratinocytes

Utilizing miRNA target prediction tools, namely Targetscan ^17^ and RNA22 ^18^, we have identified four potential binding sites for hsa-miR-129-5p (miR-129) on circASH1L (**Fig. 3A and S2A**). To experimentally validate the interaction between miR-129 and circASH1L, luciferase reporter assays were conducted in primary keratinocytes isolated from adult human skin (HEKa) and the HEK293 cell line. Overexpression of miR-129 by transfecting its mimics significantly reduced luciferase activity from the reporter construct containing the wild-type circASH1L sequence. Importantly, this effect was nullified by mutating all predicted miR-129 binding sites in the reporter construct, highlighting the necessity of these binding motifs for mediating miR-129 binding to circASH1L (**Fig. 3B and S2B**). Furthermore, we conducted a biotin pull-down assay by transfecting keratinocytes with biotin-labeled miR-129. This assay revealed an 8.7-fold higher amount of circASH1L pulled down compared to biotin-labeled negative control oligos, providing additional confirmation of the circASH1L-miR-129 interaction (**Fig. 3C**). Intriguingly, despite the presence of miR-129 binding sequences in the ASH1L mRNA, no significant amount of linear ASH1L was pulled down by miR-129 (**Fig. 3C**). This observation suggests the existence of additional mechanism(s), such as protein translation, preventing the binding of miR-129 to the coding region of ASH1L mRNA.

Furthermore, in small RNA-seq and qRT-PCR analysis of human wound samples (**Fig. 1A**), we demonstrated the upregulation of miR-129 in epidermal keratinocytes during wound healing (**Fig. 3D, 3E, and S2C**). Additionally, the expression levels of miR-129-5p were positively correlated with circASH1L in wound-edge keratinocytes in human wounds (**Fig. 3F**). Subsequently, we investigated whether circASH1L could influence miR-129 levels or *vice versa*. For this purpose, we transfected human primary keratinocytes with siRNAs targeting the BSJ of circASH1L or a circASH1L overexpression (oe) plasmid, effectively modulating circASH1L levels (**Fig. 3G**). Our findings revealed that circASH1L silencing slightly reduced the endogenous levels of miR-129, while overexpression of circASH1L significantly (P=0.03) increased miR-129 levels by 2.4-fold (**Fig. 3H**). Furthermore, we manipulated miR-129 levels in keratinocytes by transfecting its mimics or inhibitors, yet these changes did not alter circASH1L levels, as confirmed by qRT-PCR analysis (**Fig. 3I and 3J**). These results suggest that circASH1L can enhance miR-129 levels, but the expression of this circRNA is not regulated by miR-129. This discovery contrasts with the commonly reported miRNA sponging model, where circRNAs frequently lead to a decrease in the levels of their interacting miRNAs ^22^. Importantly, neither circASH1L nor miR-129 expression changes had an impact on ASH1L levels, emphasizing the independence of circASH1L’s function from its linear counterpart (**Fig. S2D and S2E**).

### CircASH1L-miR-129 axis promotes keratinocyte migration and proliferation

Given the elevated levels of both circASH1L and miR-129 in epidermal cells during wound healing, we investigated whether they play any functional roles in keratinocytes. To address this, we conducted microarray analysis in human primary keratinocytes transfected with miR-129 mimics or circASH1L siRNAs (n=3). Following miR-129 overexpression, 168 DE genes (|Fold Change| > 1.5, P < 0.05) were identified (**Table S2**), with five out of the top ten gene ontology (GO) biological processes (BP) associated with cell proliferation and migration (**Fig. 4A and Table S3**). Similarly, circASH1L silencing resulted in 960 DE genes (|Fold Change| > 1.5, P < 0.05) (**Table S4**), with seven out of the top ten GO terms related to these BP (**Fig. 4B and Table S5**). Furthermore, gene set enrichment analysis (GSEA) indicated the upregulation of genes associated with epithelial cell migration and proliferation following miR-129 overexpression (**Fig. S3A and S3B**).

**Figure 4.**
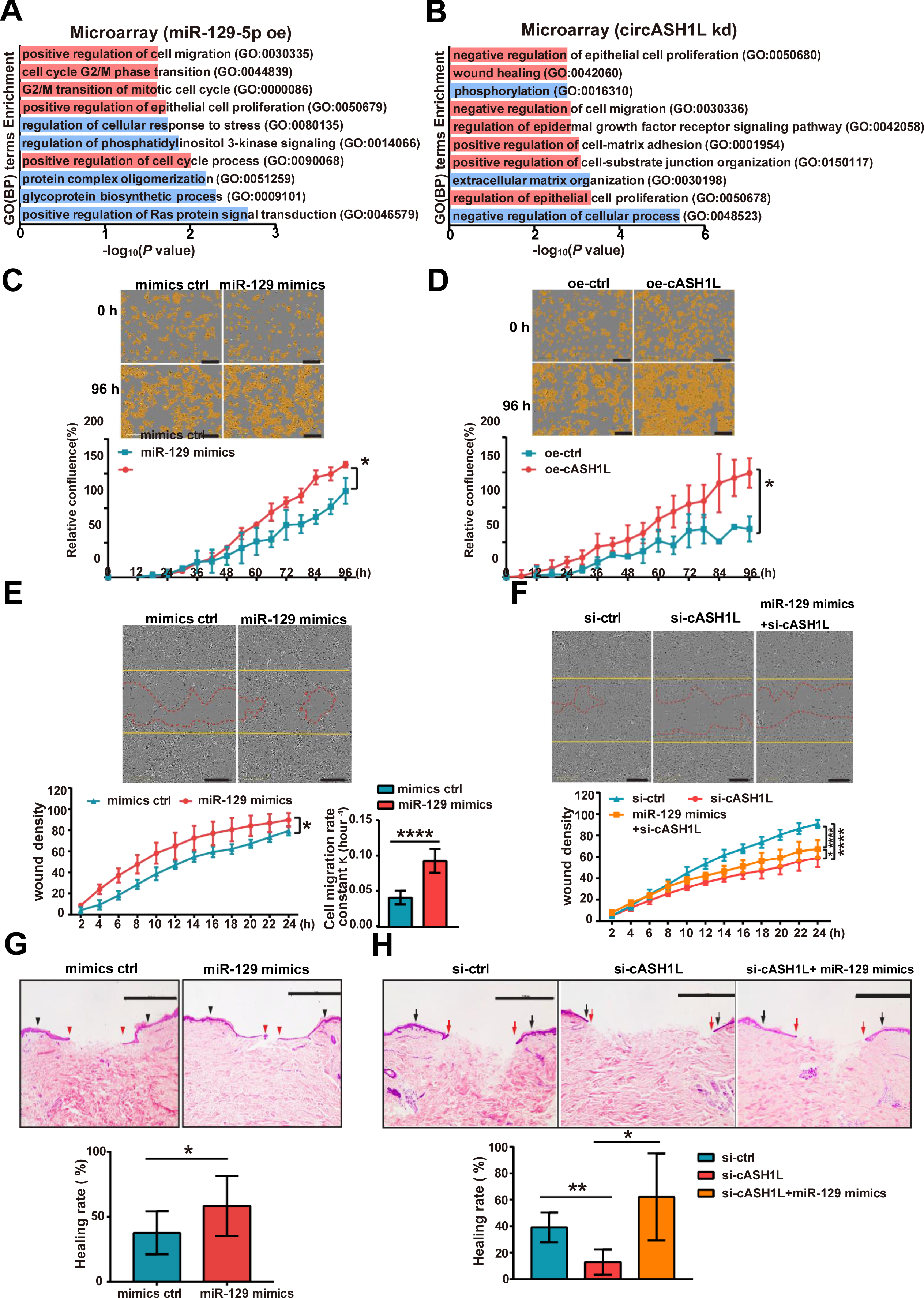
CircASH1L-miR-129 axis promotes keratinocyte migration and proliferation. Gene Ontology analysis of biological processes (BP) enriched in DE genes in HEKa cells with miR-129-5p overexpression (A) or circASH1L knockdown (B). Proliferation assays were conducted in HEKa cells with miR-129 (C) or circASH1L overexpression (D) (n=4). Scale bar = 300 μm. Cell confluence was measured at 6-hour intervals for a total duration of 96 hours. Scratch wound assays were performed in HEKa cells transfected with miR-129 mimics or control oligos (mimics ctrl) (E), and in HEKa cells transfected with control siRNA (si-ctrl), circASH1L siRNA (si-cASH1L), or co-transfected with miR-129 mimics and si-cASH1L (F) (n=4). Yellow and red lines indicate the wound-edges at 0 and 24hours, respectively. Scale bar = 300 μm. (G and H) Hematoxylin and eosin staining of human *ex vivo* wounds with topical application of miR-129 mimics and si-cASH1L on day 0 and day 3 (n=3-10). Black arrows: the initial wound edges. Red arrows: newly formed epidermis in the front (day 6). Scale bar = 1000 μm. Wound healing (%) = (1 - distance between red arrows / distance between black arrows) x100%. Data are presented as means ± SD; \**P* < 0.05; \*\**P* < 0.01; \*\*\*\**P* < 0.0001; Two-way ANOVA and multiple comparisons test (C, D, E, F); Unpaired Student’s t-test (E, G); One-way ANOVA and multiple comparisons test (H).

These *in silico* findings prompted us to investigate the impact of circASH1L and miR-129 on keratinocyte proliferation and migration. Using IncuCyte™ live-cell imaging, we observed that overexpression of either circASH1L or miR-129 significantly enhanced cell growth (**Fig. 4C and 4D**). In a scratch wound assay, keratinocytes with circASH1L or miR-129 overexpression exhibited accelerated migration, while inhibition of either circASH1L or miR-129 reduced cell migration (**Fig. 4E, F, S3C, and S3D**). Intriguingly, miR-129 overexpression improved the migration of keratinocytes with circASH1L silencing (**Fig. 4F**).

To further assess the relevance of circASH1L and miR-129 in human skin wound healing, we utilized a human *ex vivo* wound model (**Fig. S3E**). In this model, miR-129 mimics or circASH1L siRNAs or both were topically applied to partial-thickness wounds created on surgically discarded human skin immediately after injury and three days later. On day six, the wounds were collected for histological analysis. We observed that overexpression of miR-129 accelerated wound re-epithelialization (**Fig. 4G**), whereas silencing of circASH1L delayed re-epithelialization, and this effect was reversed by adding miR-129 mimics (**Fig. 4H**). Based on these results, we concluded that circASH1L and miR-129 promote keratinocyte migration and proliferation and are required for human wound re-epithelialization.

### CircASH1L downregulates the expression of miR-129’s target genes

In the commonly reported circRNA sponging miRNA model ^3,22^, a decrease in circRNA expression leads to the release of its sponged miRNAs, subsequently suppressing the expression of miRNA target genes. Unexpectedly, upon silencing circASH1L, we observed the upregulation of most miR-129’s target mRNAs, as demonstrated by GSEA of our microarray data (**Fig. 5A**). A heatmap revealed significantly increased expression (fold change > 1.5, P < 0.05) of several miR-129 targets in keratinocytes with circASH1L silencing (**Fig. 5B**).

**Figure 5.**
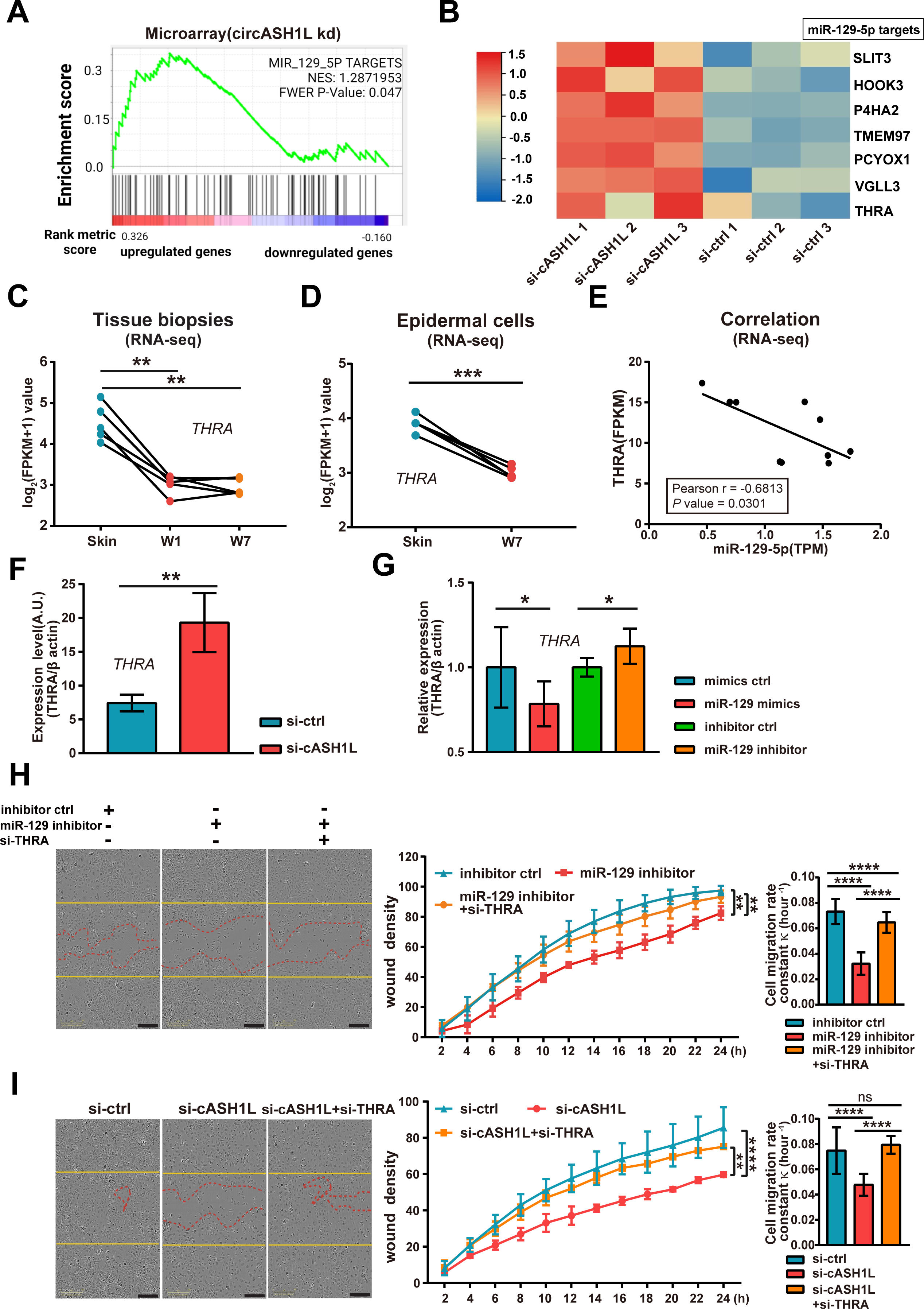
CircASH1L downregulates the expression of miR-129’s target genes. (A) Gene set enrichment analysis (GSEA) shows that miR-129 targets are enriched in the genes upregulated in HEKa with circASH1L knockdown (kd). (B) Heatmap depicts the expression of seven miR-129-5p targets in HEKa with circASH1L kd, as shown in microarray analysis (n=3). (C and D) RNA sequencing data showing THRA expression in the skin, day-1, and day-7 wound tissues (n=5 donors) (C) and in the skin and day-7 wound epidermal cells (n=5 donors) (D). (E) Correlation analysis of THRA and miR-129-5p levels in the skin and day-7 wound epidermal cells (n=10). qRT-PCR analysis of THRA expression in HEKa cells transfected with circASH1L siRNA (si-cASH1L) or control siRNA (si-ctrl) (F), and in HEKa cells transfected with miR-129 mimics, miR-129 inhibitors, or their negative controls (n=4) (G). (H) Scratch wound assays of HEKa cells transfected with inhibitor control (ctrl), or miR-129 inhibitor, or co-transfected with miR-129 inhibitor and THRA siRNA (si-THRA) (n=5). (I) Scratch wound assays of HEKa cells transfected with si-ctrl, si-cASH1L, or co-transfected with si-cASH1L and si-THRA (n=4). Yellow and red lines indicate the wound-edges at 0 and 24hours, respectively. Scale bar = 300 μm. Data are presented as individual values or means ± SD; \**P* < 0.05; \*\**P* < 0.01; \*\*\**P* < 0.001; \*\*\*\**P* < 0.0001; One-way ANOVA and multiple comparisons test (C, H, I); Paired Student’s t-test (D); Pearson’s correlation test (E); Unpaired Student’s t-test (F, G); Two-way ANOVA and multiple comparisons test (H, I).

Investigating one of these miR-129 targets, Thyroid Hormone Receptor Alpha (THRA) (**Fig. S4A**), we found its expression significantly decreased in epidermal cells during wound healing (**Fig. 5C and 5D**). Notably, THRA expression negatively correlated (Pearson coefficient r = -0.68, P = 0.03) with miR-129 levels in human wound-edge keratinocytes, supporting the notion that miR-129 targets THRA (**Fig. 5E**). Consistent with this, both circASH1L silencing and miR-129 inhibition increased THRA expression, while overexpression of miR-129 and circASH1L reduced THRA levels in human keratinocytes (**Fig. 5F, 5G, and S4B**).

Functionally, scratch wound assays revealed that silencing THRA expression enhanced keratinocyte migration, identifying it as a suppressor of cell motility (**Fig. S4C**). Intriguingly, THRA silencing could rescue the delayed migration of keratinocytes with either miR-129 inhibition or circASH1L silencing, indicating that THRA is a key mediator of circASH1L and miR-129 function (**Fig. 5H and 5I**).

Collectively, our data suggest that, different from the miRNA sponging model, circASH1L enhances miR-129 levels, subsequently suppressing the expression of its targets, such as THRA, a negative regulator of cell motility. Therefore, the elevated expression of both circASH1L and miR-129 is crucial for promoting keratinocyte migration and wound re-epithelialization.’

### CircASH1L enhances miR-129 stability by competing with NR6A1-triggered TDMD

We next investigated the mechanism by which circASH1L increases miR-129 levels in keratinocytes. Initially, we examined whether circASH1L could impact miR-129 biogenesis. Mature miR-129 is derived from two primary precursors, pri-miR-129-1 and pri-miR-129-2. Neither silencing nor overexpression of circASH1L altered the expression levels of miR-129 precursors, indicating that circASH1L does not influence miR-129 expression at the transcriptional level (**Fig. S5A and S5B**). Additionally, we observed no significant change in the expression of these miR-129 precursors during human skin wound healing, as indicated by qRT-PCR analysis of human skin and wound biopsies (**Fig. S5C and S5D**).

Next, we explored whether circASH1L might interfere with miR-129 degradation. Treating human primary keratinocytes with actinomycin-D (5 μg/mL), which inhibits total cellular transcription ^23^, revealed that miR-129 and circASH1L exhibited similar stability, significantly higher than that of linear ASH1L mRNA (**Fig. S5E**). Interestingly, overexpression of circASH1L further enhanced miR-129 stability, while circASH1L silencing led to accelerated degradation of miR-129 (**Fig. 6A and 6B**). Thus, we concluded that circASH1L primarily regulates miR-129 cellular levels by reducing the degradation of this miRNA.

**Figure 6.**
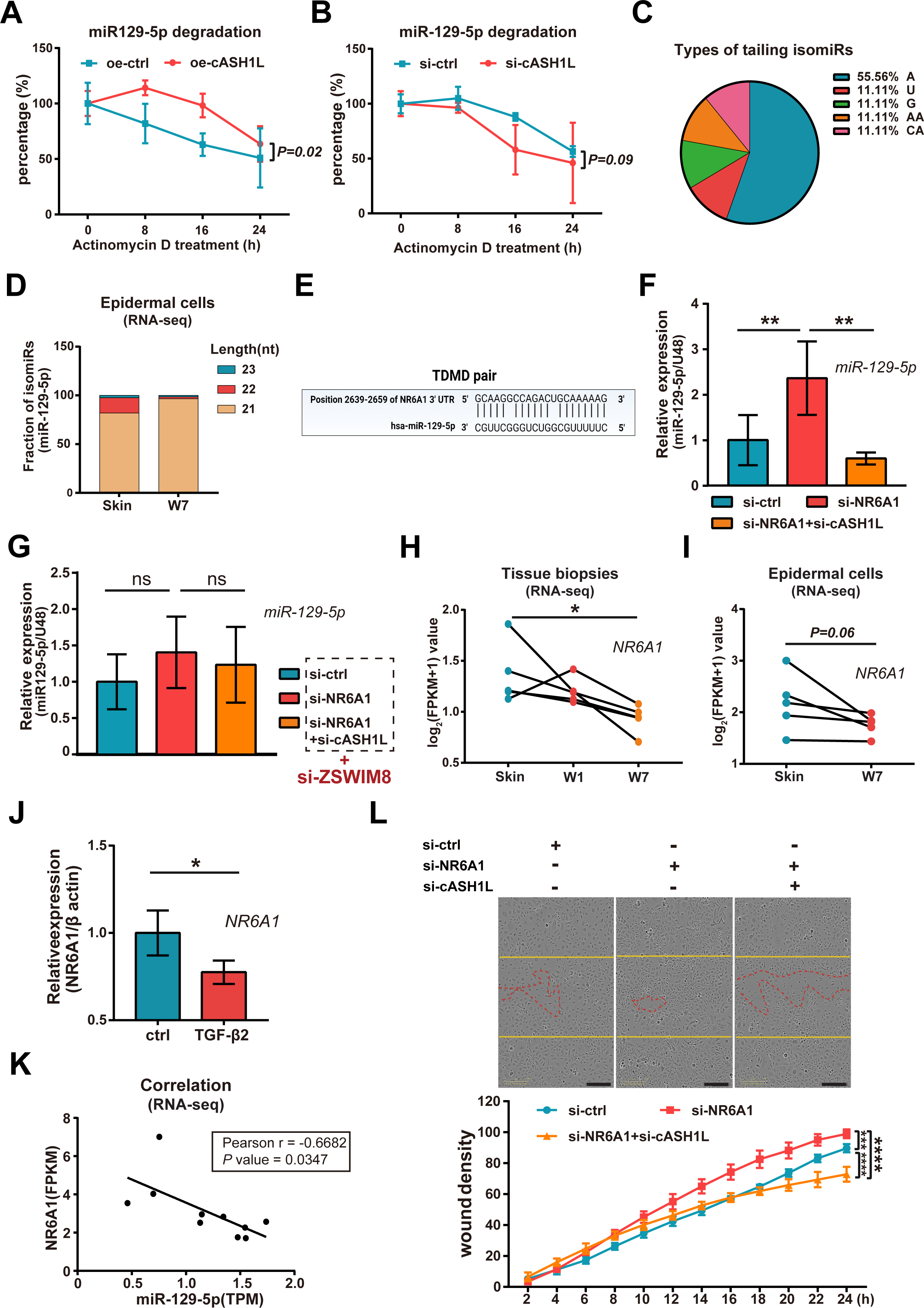
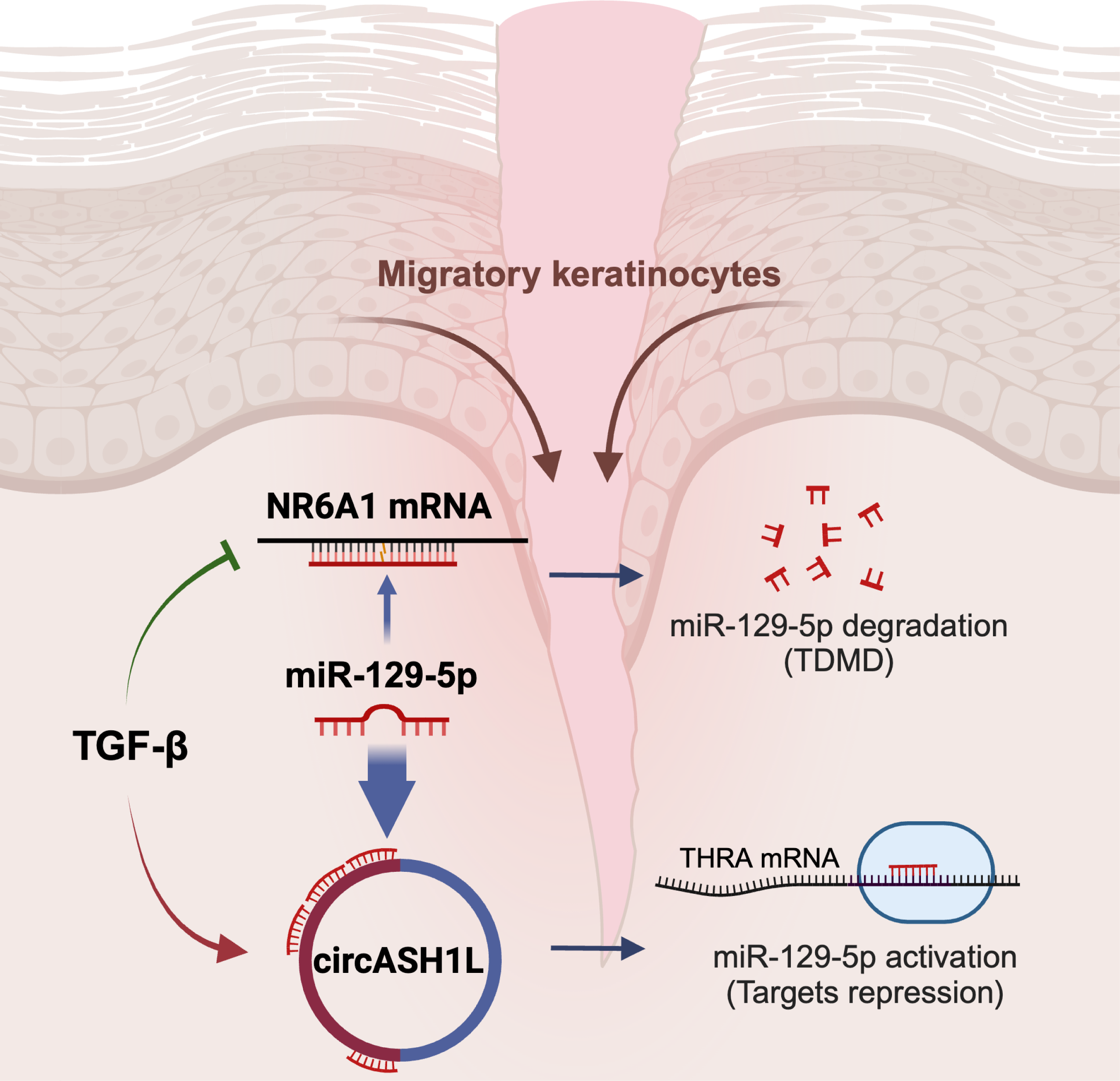
CircASH1L enhances miR-129 stability by competing with NR6A1-triggered TDMD. qRT-PCR analysis of miR-129-5p in HEKa cells with circASH1L overexpression (A) or knockdown (B) and treated with actinomycin D for 0-24hours (n=4). Analysis of miR-129-5p isoforms in the small RNA-seq data of human skin and day-7 wound (W7) epidermal cells: pie chart depicts the types of tailing isomiRs (C); bar chart shows the changes of tailing isomiR fractions in wounds compared to the skin (D). (E) Illustration of the miR-129-5p binding site at NR6A1 3’UTR. qRT-PCR of miR-129-5p in HEKa cells transfected with si-NR6A1 or co-transfected with si-NR6A1 and si-cASH1L, and not with (F) or with ZSWIM8 siRNA (G) (n=3). RNA sequencing data showing NR6A1 expression in the skin, day-1, and day-7 wound tissues (n=5 donors) (H) and in the skin and day-7 wound epidermal cells (I) (n=5 donors). (J) qRT-PCR of of NR6A1 in HEKa cells treated with TGF-β2 (n=3). (K) Correlation analysis of NR6A1 and miR-129-5p in skin and wound epidermal cells (n=10). (L) Scratch wound assay of HEKa cells transfected with si-ctrl, si-NR6A1, or co-transfected with si-NR6A1 and si-cASH1L (n=4). Scale bar = 300 μm. Data are presented as means ± SEM (A, B), or means ± SD (F, G, J, L), or individual values (H, I, K); \**P* < 0.05; \*\**P* < 0.01; \*\*\*\**P* < 0.0001; Two-way ANOVA and multiple comparisons test (A, B, L); One-way ANOVA and multiple comparisons test (F, G, H); Paired Student’s t-test (I); Unpaired Student’s t-test (J); Pearson’s correlation test (K).

As a major mechanism regulating turnover of specific animal miRNAs, target-directed miRNA degradation (TDMD) often leads to miRNA tailing or trimming before complete degradation, generating miRNA isoforms (isomiRs) ^4,24,25^. Small RNA-seq analysis of human skin and Day 7 wound-edge epidermal cells identified five types of tailing isomiRs for miR-129, carrying one or two additional nucleotides at the 3’-end of miR-129, with adenine (A) being the most common type (**Fig. 6C**). No trimming miR-129 isomiRs were observed. In keratinocytes from intact skin, isomiRs constituted approximately 20% of total miR-129, whereas this fraction decreased to 4% in wound-edge keratinocytes, indicating reduced miR-129 degradation during wound repair (**Fig. 6D and S5F**).

We sought TDMD-inducing targets for miR-129, expecting these targets to pair with miR-129 not only at the 5′ end (seed region) but also at the 3′ end of the miRNA and carry central mismatches ^26^. One miR-129 binding site at the 3′ untranslated regions (3′ UTR) of NR6A1 mRNA (Nuclear receptor subfamily 6 group A member 1) met these criteria (**Fig. 6E**). Overexpression or inhibition of miR-129 did not change NR6A1 expression due to central mismatches preventing AGO-mediated cleavage of the target NR6A1 mRNA (**Fig. S6A**) ^4^. However, silencing NR6A1 expression significantly elevated miR-129 levels in human keratinocytes, and co-silencing of circASH1L counteracted the effects of NR6A1 silencing on miR-129 (**Fig. 6F and S6B**). Interestingly, in keratinocytes lacking ZSWIM8, a ubiquitin ligase mediating TDMD, silencing NR6A1 did not alter miR-129 levels, supporting miR-129 as a *bona fide* substrate of TDMD triggered by NR6A1 (**Fig. 6G and S6C**).

Furthermore, we noted a decline in NR6A1 mRNA expression in epidermal cells during wound healing (**Fig. 6H and 6I**), potentially attributed to injury-induced TGF-β signaling inhibiting NR6A1 expression (**Fig. 6J**) ^27^. Additionally, NR6A1 and miR-129 levels were negatively correlated in human skin and wound keratinocytes, supporting NR6A1-induced miR-129 degradation *in vivo* (**Fig. 6K**). Functionally, silencing NR6A1 accelerated keratinocyte migration, and this effect could be reversed by co-silencing circASH1L (**Fig. 6L**). Moreover, silencing NR6A1 also promoted keratinocyte growth (**Fig. S6D**).

In concert, our findings propose a balanced model governing miR-129 turnover: circASH1L binds to miR-129, shielding it from degradation, while NR6A1 mRNA triggers miR-129 degradation via TDMD. This equilibrium can be influenced, as exemplified by TGF-β signaling, which enhances circASH1L levels while diminishing NR6A1 levels during human skin wound healing. Consequently, this leads to an elevation of miR-129 cellular levels. The heightened miR-129, in turn, facilitates keratinocyte migration and wound re-epithelialization by suppressing its targets, such as THRA. Therefore, our study presents compelling evidence that circular RNA engages with the endogenous TDMD mechanism, orchestrating the turnover of a miRNA pivotal for keratinocyte functions during human skin wound healing.

## Discussion

MiRNAs play a crucial role in regulating gene expression, influencing various physiological and pathological processes such as wound healing, making them potential therapeutic entities^11^. While substantial knowledge exists about miRNA biogenesis and biological activities, understanding of miRNA turnover and its impact on human health and disease remains limited ^4^. Recently, target-directed miRNA degradation (TDMD) has been recognized as a major mechanism regulating the turnover of animal miRNAs ^4^. Over the past few years, numerous naturally occurring examples of TDMD have surfaced, spanning both endogenous and viral transcripts, impacting biological processes such as infection, cancer, embryonic development, and animal behavior ^24,28-33^. Our study contributes to this understanding by providing compelling evidence that TDMD operates in human epidermal keratinocytes and plays a functional role in human skin wound healing. We identified NR6A1 mRNA as a novel endogenous trigger for miR-129 degradation, adding to the growing list of mRNAs with non-coding functions ^34^. Our findings underscore the importance of endogenous TDMD in human physiology, with the likelihood of more examples awaiting discovery.

Adding to the novelty of our study, we unveil a significant interaction between circRNAs and the TDMD mechanism, akin to the case reported in mouse brain where circRNA Cdr1as protects miR-7-5p from TDMD triggered by the lncRNA Cyrano ^8,9^. Intriguingly, recent findings by Wightman F.F. *et al.* highlight that the miRNA-stabilizing effects of Cdr1as depend on its circular topology, as the expression of an artificially linear version of Cdr1as induces TDMD^10^. The inability of Cdr1as to drive the TDMD of miR-7, coupled with its capacity to compete with miR-7 for the TDMD trigger Cyrano, ultimately leads to the stabilization of miR-7^10^. In alignment with this paradigm, our investigation demonstrates that circASH1L binds to miR-129 (**Fig. 3B and 3C**); however, it does not alter miR-129 levels in keratinocytes lacking the TDMD trigger NR6A1 or the essential ubiquitin ligase ZSWIM8 (**Fig. 6F and 6G**). When the TDMD mechanism is intact, silencing or overexpressing circASH1L diminishes or augments miR-129 stability, respectively (**Fig. 6A and 6B**). This suggests that circASH1L can compete with NR6A1 for miR-129 binding, thereby shielding miR-129 from NR6A1-induced TDMD in human keratinocytes.

Similarly to the scenario involving miR-7, where its stabilization by Cdr1as led to greater target repression ^8,9^, the decreased miR-129 abundance due to the loss of circASH1L also resulted in the release of the expression of miR-129’s targets, such as THRA, in human keratinocytes. Our study underscores that circASH1L enhances miR-129’s silencing activity, contrasting with the widely reported sponging model, where circRNAs typically bind to miRNAs, preventing their interaction with targets and thus reducing miRNA functions ^3^. This reveals that the interaction between circRNAs and miRNAs can yield different outcomes in the canonical miRNA silencing activity^10^. Building upon just two existing cases—Cdr1as with miR-7 and circASH1L with miR-129—we speculate whether the miRNA protective role of circRNAs may be exclusive to miRNAs subjected to TDMD. This prompts the need for further studies dissecting the impact of circRNAs on miRNA turnover. A more profound molecular understanding of how circASH1L shields miR-129 from TDMD triggers while facilitating its interaction with targets is essential.

Notably, the pivotal role of miR-129 in wound healing had been recognized in diabetic wounds before our study ^35^. Its expression is diminished in human diabetic ulcers compared to normal skin, and the application of miR-129 agomirs in diabetic animals has been shown to accelerate keratinocyte migration and enhance wound healing ^35^. Consistent with the pro-migratory and pro-proliferative functions of miR-129, our study revealed an upregulation of this miRNA in human wound-edge keratinocytes during wound repair. This surge in miR-129 abundance primarily occurred at the post-transcriptional level, attributable to the heightened expression of its protective counterpart, circASH1L, and the reduced expression of its TDMD-triggering factor, NR6A1. Intriguingly, both circASH1L upregulation and NR6A1 downregulation in keratinocytes are induced by TGF-β signaling—a well-known growth factor produced by platelets, macrophages, and keratinocytes in the wounds and plays diverse roles in wound healing, particularly in regulating keratinocyte migration and proliferation during re-epithelization^27^. Moreover, TGF-β has been shown to modulate the maturation of miRNAs possessing a SMAD-binding element-like sequence in their primary transcripts. This regulation involves the association of Smads with the Drosha microprocessor complex, resulting in the upregulation of pre-miRNAs and matured miRNAs, but not of pri-miRNAs^36,37^. Consequently, the TGF-β signaling pathway establishes a multifaceted crosstalk with the miRNA machinery at various levels ^38^.

In summary, our study sheds light on the physiological importance of regulated miRNA degradation in human skin wound healing. We uncover a novel function of circRNAs, acting as shields for their interacting miRNAs from TDMD, thereby enhancing miRNA activity. This endogenous mechanism provides an alternative to the well-known miRNA sponging model, allowing for customization of specific miRNA abundance. As synthetic circRNA sponges are considered potential therapeutic strategies, our findings emphasize the need for careful examination of their impact on interacting miRNAs, including turnover and activity ^39^. Further exploration of when, where, and how circRNAs exert their miRNA-protective role will deepen our understanding of the intricate network of noncoding regulatory RNAs in both health and disease.

## Supporting information

Table S1 Differentially expressed circRNAs in human wound tissues and epidermal cells

Table S2 DE genes in keratinocytes with miR-129 overexpression

Table S3 GO analysis of DE genes in keratinocytes with miR-129 overexpression

Table S4 DE genes in keratinocytes with circASH1L silencing

Table S5 GO analysis of DE genes in keratinocytes with circASH1L silencing

## Acknowledgments

We thank all the tissue donors participating in this study. We thank the Microarray core facility at Novum, BEA, which is supported by the board of research at KI and the research committee at the Karolinska hospital. This work was supported by Swedish Research Council (Vetenskapsrådet) (2020-01400), Ragnar Söderbergs Foundation (M31/15), Welander and Finsens Foundation (Hudfonden), Ming Wai Lau Centre for Reparative Medicine, LEO Foundation, Cancerfonden, Karolinska Institutet, and China Scholarship Council (202006230148). The computations/data handling was enabled by resources in projects of sens2020010 and naiss2023-22-935 provided by the Swedish National Infrastructure for Computing (SNIC) at UPPMAX, partially funded by the Swedish Research Council through grant agreement no. 2018-05973. Schematic cartoons in Fig. 1A and the graphical abstract were created with BioRender.com.

## Author contributions

N.X.L. and Q.W. conceptualized the study. Q.W., G.N., M.T., J.G., Z.L., X.B., L.Z., and M.P. conceived and performed experiments. Q.W., N.X.L, M.T., and Z.L. analyzed the data. N.X.L., G.N., Z.L., and Q.W. provided funding for the study. Q.W., X.B., N.X.L., and D.L. were responsible for study execution. N.X.L., P.S., D.L. supervised the study. N.X.L and Q.W. wrote the manuscript. All authors read and approved the final version of the manuscript.

## STAR★Methods

## KEY RESOURCES TABLE

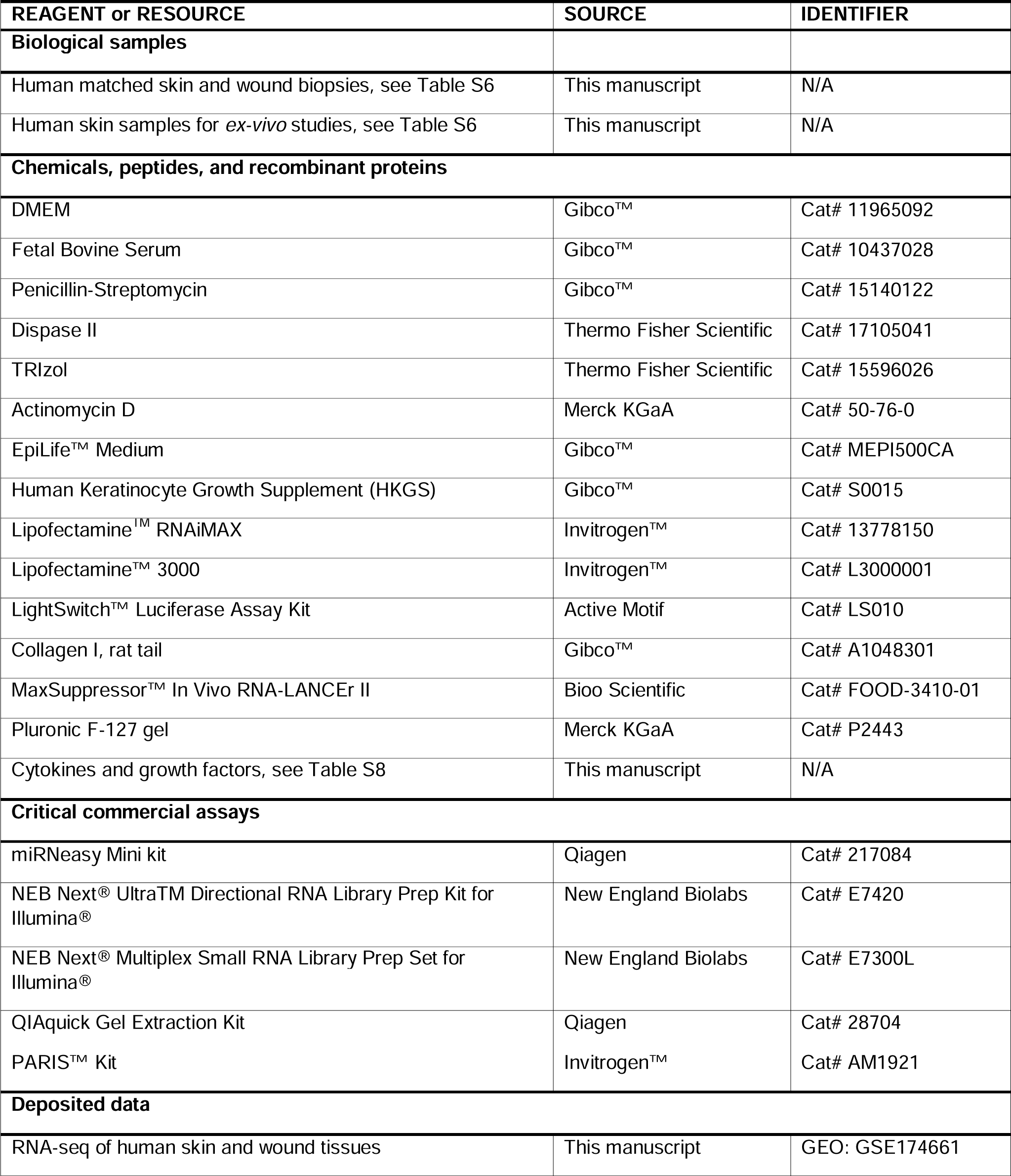
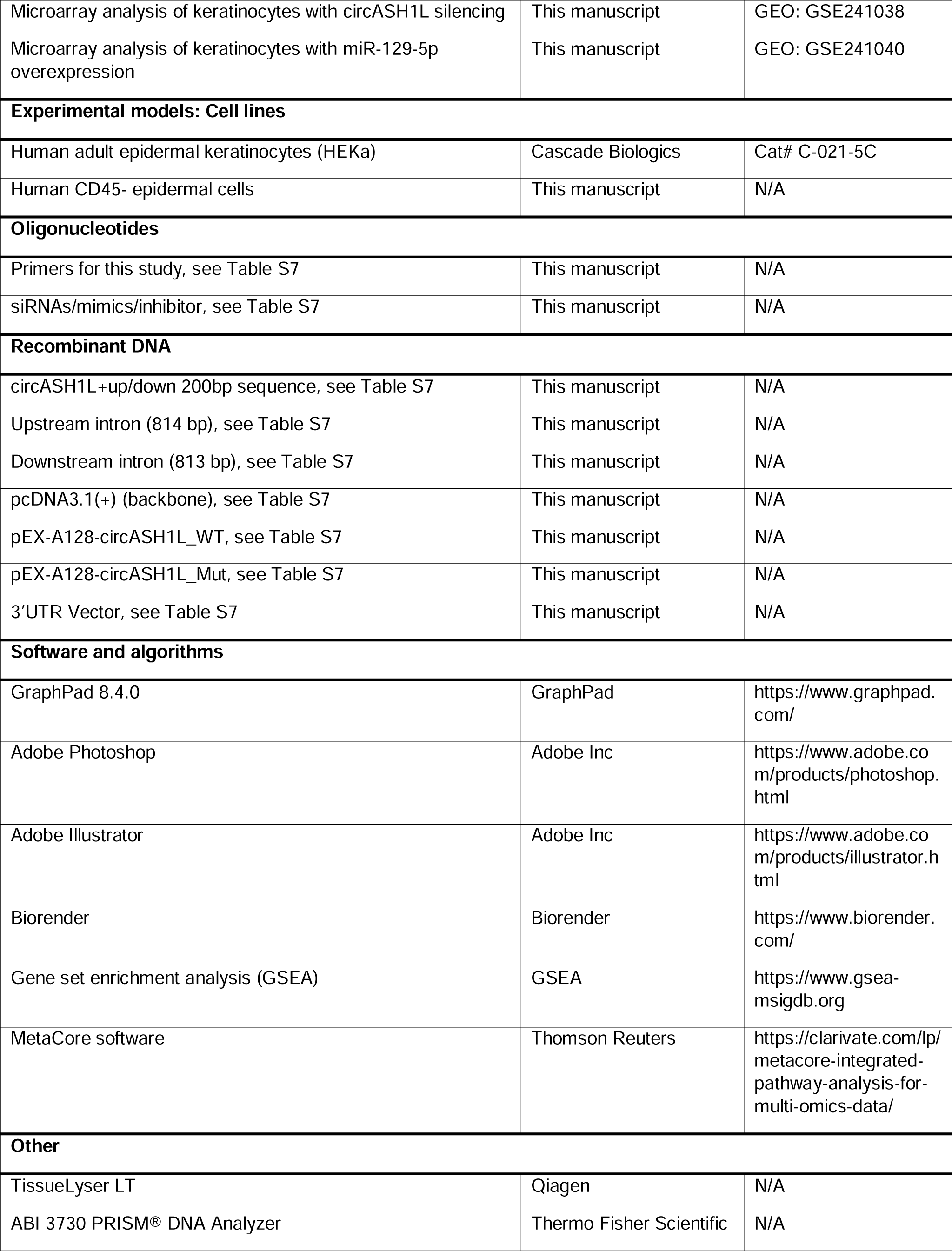
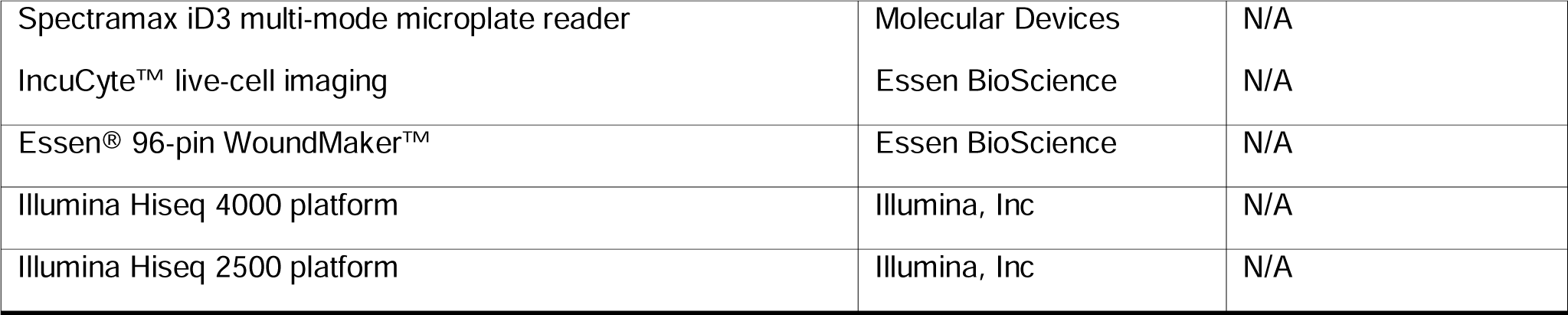

## RESOURCE AVAILABILITY

### Lead contact

Further information and requests for resources and reagents should be directed to and will be fulfilled by the lead contact, Ning Xu Landén.

### Materials availability

This study did not generate new unique reagents.

### Data and code availability

Raw data of RNA-seq and microarray in this study have been deposited to NCBI’s Gene Expression Omnibus (GEO) database under the accession numbers GSE174661 (RNA-seq of human skin and wound tissues), GSE241038 (token: etufciykfjazvqt, microarray analysis of keratinocytes with circASH1L silencing), and GSE241040 (token: uvexwawqdpohban, microarray analysis of keratinocytes with miR-129-5p overexpression).

Any additional information required to reanalyze the data reported in this paper is available from the lead contact upon request.

## EXPERIMENTAL MODEL AND STUDY PARTICIPANT DETAILS

### Human skin wound specimens

Fifteen healthy volunteers provided matched skin and wound biopsies, with donor details outlined in **Table S6**. Each volunteer had two 4-mm excisional wounds created on their skin for study, with the removed skin serving as an intact baseline. Wound margins were then harvested with a 6-mm biopsy punch at one and seven days after injury. Local anesthesia with lidocaine was administered during these procedures. Additionally, *ex-vivo* wound healing studies were performed on human skin discarded from surgeries (**Table S6**). All participants gave their informed consent in writing for the collection and utilization of their samples. The research protocol was approved by the Stockholm Regional Ethics Committee and was conducted in accordance with the Declaration of Helsinki’s guidelines.

### Human ex vivo wound model

Human skin for *ex vivo* wound models was obtained from plastic surgeries. The wounds were created using a 2 mm biopsy punch on the epidermal side of the skin. The wounds were then excised using a 6 mm biopsy punch and cultured in 24-well plates with Dulbecco’s Modified Eagle Medium (DMEM), 10% fetal bovine serum (FBS), and antibiotics (penicillin 50 U/l and streptomycin 50 mg/ml, ThermoFisher Scientific). MaxSuppressor In Vivo RNA-LANCEr II (Bioo Scientific, Austin, TX) was mixed with 0.1 nmol siRNA targeting circASH1L or a scrambled siRNA or miRIDIAN miR-129 mimics or control mimics in a volume of 5 µl per wound. The siRNA-lipid complexes were mixed 1:2 (volumes) in 30% pluronic F-127 gel (Sigma-Aldrich). 15 µl mixture was topically applied on the wounds immediately after injury and 3 days later. Wound samples were collected on day 6 and used for histological analysis.

### Magnetic cell separation of CD45-epidermal cells from human skin and wound biopsies

Following PBS wash, skin and wound tissues were incubated in dispase II (5 U/mL, ThermoFisher Scientific, Carlsbad, CA) at 4°C overnight, leading to the separation of the epidermis and dermis. The epidermis was then sectioned and subjected to digestion in 0.025% Trypsin/EDTA Solution at 37°C for 15 minutes. The isolation of CD45- and CD45+ cells was accomplished using CD45 Microbeads and MACS MS magnetic columns, following the manufacturer’s instructions (Miltenyi Biotec, Bergisch Gladbach, Germany).

### Cell lines

Human adult epidermal keratinocytes (HEKa) (Cascade Biologics, Portland, OR) were cultured in EpiLife medium (ThermoFisher Scientific, Carlsbad, CA) supplemented with 10% Human Keratinocyte Growth Supplement (HKGS) as well as 1% penicillin/streptomycin. Cells were normally incubated at 37°C in 5% CO_2_ (ThermoFisher Scientific) and the third passages were used for the experiments.

## METHOD DETAILS

### RNA extraction

Tissue biopsies were homogenized using TissueLyser LT (Qiagen) before RNA extraction. Total RNA was extracted from human tissue and cells using the miRNeasy Mini kit (Qiagen, Hilden, Germany) or TRIzol reagent (Thermo Fisher Scientific).

### RNA sequencing

For long RNA sequencing, the ribosomal RNA (rRNA) was removed using the Epicentre Ribo-zero® rRNA Removal Kit (Epicentre, Road Madison, WI). Subsequently, RNA-seq libraries were constructed using the NEB Next® UltraTM Directional RNA Library Prep Kit for Illumina® (New England Biolabs, Ipswich, MA). Finally, the libraries were sequenced on the Illumina Hiseq 4000 platform (Illumina, Inc., San Diego, CA) using 150 bp paired-end reads. Raw reads were firstly filtered by the Trimmomatic v 0.36 package. Clean reads were then mapped to human reference genome (GRCh38.p12) with the GENCODE genes annotation (version 31) using STAR v2.7.1. Gene expression of mRNAs was obtained by counting specifically mapped fragments to exons using featureCount package ^40^. The unmapped chimeric reads were used for circRNA identification using CIRCexplorer2 and DCC. CircRNAs were filtered by at least two back-spliced junction reads in a minimum of two samples. CircRNAs were annotated based on chromosomal location as well as the overlap with three circRNA databases: circAtlas v2, circBase, and CIRCpedia v2.

For small RNA sequencing, the libraries were firstly constructed using the NEB Next® Multiplex Small RNA Library Prep Set for Illumina® (NEB). Total RNA was then ligated to adaptors at the 3’ end by NEB 3’ SR adaptor and 5’ end by T4 RNA ligase, respectively, followed by reverse transcription into cDNA using M-MuLV Reverse Transcriptase. PCR amplification was carried out using SR primers for Illumina and index primers. The PCR products were then purified and DNA fragments spanning from 140 to 160 bp were recovered and quantified by DNA High Sensitivity Chips on the Agilent Bioanalyzer. The libraries were sequenced on an Illumina Hiseq 2500 platform (Illumina, Inc.) using single-end 50 bp reads.

### Quantitative real-time reverse transcription polymerase chain reaction (qRT-PCR)

qRT-PCR were conducted as previously described ^41^. Gene expression was examined by SybrGreen expression assays (ThermoFisher Scientific) or TaqMan expression assays (ThermoFisher Scientific) and normalized according to the values of the housekeeping genes: RNU48, GAPDH, β actin, or 18S. The sequences for all the primers and probes used in this study were listed in **Table S7**.

### RNase R treatment, DNA electrophoresis, and Sanger sequencing

To confirm circASH1L’s circular nature, human keratinocyte-isolated RNA underwent treatment with RNase R at 37°C for 30 or 60 minutes, followed by inactivation at 70°C for 10 minutes. RT-PCR with divergent primers spanning the circRNA-forming back-splice junction was conducted. Subsequent analysis of PCR products occurred through DNA electrophoresis using E-Gel Agarose Gels with SYBR Safe DNA Gel Stain, 2% (Invitrogen, Waltham, MA). Bands of expected sizes were excised, and DNA purification was carried out using the QIAquick Gel Extraction Kit (Qiagen). The purified DNA underwent Sanger sequencing using the ABI 3730 PRISM DNA Analyzer at the KIgene core facility at Karolinska Institutet (Stockholm, Sweden).

### Cell fractionation

Nuclear and cytoplasmic fractions were separated by PARIS kit following manufacturer’s instructions (ThermoFisher Scientific). RNA was extracted from these fractions by using Trizol (ThermoFisher Scientific) and qRT-PCR was performed to analyze the MALAT1, GAPDH, and circASH1L expression.

### Cell treatments

To study the mechanisms regulating circASH1L expression, we treated keratinocytes with cytokines and growth factors (**Table S8**), or PBS as a control for 24 hours, and then circASH1L expression was analyzed by qPCR.

To study the biological functions of circASH1L and miR-129, we transfected the third passage of keratinocytes at 60-70% confluence with 20nM siRNA targeting circASH1L (si-cASH1L) or siRNA controls (si-ctrl) (Dharmacon, Cambridge, UK); 20nM hsa-miR-129-5p miRIDIAN mimics (miR-129 mimics) or mimic negative control (mimics ctrl)(ThermoFisher Scientific); 20nM has-miR-129-5p miRIDIAN hairpin inhibitor (miR-129 inhibitor) or hairpin inhibitor negative control (inhibitor ctrl) (ThermoFisher Scientific) using Lipofectamine^TM^ RNAiMAX (ThermoFisher Scientific). To overexpress circASH1L, we transfected keratinocytes in 24-well plates with 500ng/well circASH1L overexpressing plasmids (oe-cASH1L) or pcDNA3.1 vector (oe-ctrl) using Lipofectamine^TM^ 3000 reagent (ThermoFisher Scientific).

To evaluate the stability of miR-129, we treated keratinocytes with with 5 μg/mL Actinomycin D (Merck KGaA, Darmstadt, Germany) multiple time points, and miR-129 levels were examined by qRT-PCR.

### Plasmid construction

CircASH1L overexpression plasmid was constructed as previously described ^42^. Briefly, full length of circASH1L together with its upstream and downstream 200bp intron fragments was subcloned into the pcDNA3.1 vector. For the PLS-cASH1L-wild plasmid, full length of circASH1L was subcloned to the 3′UTR region of pLightSwitch backbone (Active Motif, La Hulpe, Belgium). For the PLS-cASH1L-mut plasmid, full length of circASH1L with four mutant sites (564-582 bp, 737-752 bp, 779-800 bp, 800-822 bp) was subcloned into the 3′ UTR region of pLightSwitch backbone. All the plasmid sequences were confirmed by sanger sequencing.

### Luciferase reporter assay

Keratinocytes were cultured in 24-well plates and then transfected with PLS-cASH1Lwild or mutant plasmids with 20 nM miR-129-5p mimics or mimic ctrl, using the Lipofectamine^TM^ 3000 reagent (ThermoFisher Scientific). Luciferase activity was measured 24 hours post-transfection using a LightSwitch Luciferase Assay kit (Active Motif) on Spectramax iD3 multi-mode microplate reader (Molecular Devices, USA). All assays were performed in triplicate.

### RNA pull-down assay

RNA pull-down assay was conducted as previously described ^43^. Briefly, keratinocytes were transfected with biotin-labeled hsa-miR-129-5p mimics, or biotin-labeled controlled oligos (NC) (Dharmacon), or without transfection (Blank) for 48 hours. Cell lysates were prepared from the transfected cells and incubated with streptavidin magnetic beads for one hour at room temperature and three hours at 4°C. Total RNA was extracted from the beads using TRIzol reagent.

### Scratch wound assay and cell proliferation assay using IncuCyte Live cell imaging

Cell migration and proliferation were analyzed with IncuCyte™ live-cell imaging (Essen BioScience, Ann Arbor, MI). For scratch wound assay, transfected keratinocytes were seeded onto a 96-well microtiter IncuCyte® ImageLock Plate (Essen BioScience) pre-coated with rat tail Collagen type I (Gibco BRL, Gaithersburg, MD) overnight. Standardized wounds were created by Essen® 96-pin WoundMaker™ (Essen BioScience) once cells became fully confluent. The medium was then switched to basic EpiLife medium without any supplements. Cells were incubated in the IncuCyte and scanned every 2 hours for a total of 24 hours. The data were analyzed using the IncuCyte software and presented as the wound density.

For proliferation assay, transfected cells were seeded at the density of 2000 cells/well in 96-well microtiter IncuCyte® ImageLock Plate (Essen BioScience) pre-coated with rat tail Collagen type I (Gibco BRL) overnight. After 24 hours, plates were incubated in the IncuCyte and scanned every 6 hours for a total of 96 hours. The cell confluence was calculated using the IncuCyte software and displayed as the relative confluence.

### Gene expression microarray and analysis

Human primary keratinocytes were transfected with either siRNA control, or siRNA targeting circASH1L, or hsa-miR-129-5p miRIDIAN mimic, or mimic negative control for 24 hours. RNA was extracted from these transfected cells and sent to the Bioinformatics and Expression Analysis (BEA) core facility at Karolinska Institutet for Human Clariom™ S assay (ThermoFisher Scientific). Briefly, 150 ng of total RNA was used for reverse transcription followed by the GeneChip WT PLUS Reagent Kit labeling protocol. Standardized array processing procedures suggested by Affymetrix, including hybridization, fluidics processing, and scanning, were used. Expression data were interpreted according to Transcriptome Analysis Console 4.0.

Genes showing at least 1.5-fold change and with a P value less than 0.05 were considered differentially expressed. Gene set enrichment analysis (GSEA) was performed using a public software from Broad Institute ^44^. MetaCore software (Thomson Reuters, Toronto, Canada) was used for Gene Ontology (GO) analysis. Heatmaps were generated with the Multiple Experiment Viewer software.

## QUANTIFICATION AND STATISTICAL ANALYSIS

Data analysis for this study employed GraphPad 8.4.0 (GraphPad Software, San Diego, CA). Shapiro-Wilk tests were used to assess data normality and distribution. Unpaired or paired Student’s t-tests were applied for two-group comparisons, while one-way or two-way ANOVA, along with multiple comparisons tests, were conducted for comparisons involving more than two groups. Pearson’s correlation test gauged the expression pattern correlation among different subjects. Each experiment was replicated at least three times, and statistical significance was defined as P < 0.05.

## Supplemental information

Figure S1. qRT-PCR analysis of circASH1L in human skin, day 1, and day 7 wound tissues.

Figure S2. CircASH1L binds to miR-129-5p and increases its level in human wound-edge keratinocytes.

Figure S3. CircASH1L-miR-129 axis promotes keratinocyte migration and proliferation. Figure S4. miR-129-5p promotes keratinocyte migration by inhibiting THRA.

Figure S5. CircASH1L enhances miR-129 stability.

Figure S6. NR6A1 is a TDMD trigger and suppresses keratinocyte growth.

Table S1. Differentially expressed circRNAs in human wound tissues and epidermal cells.

Table S2. DE genes in keratinocytes with miR-129 overexpression.

Table S3. GO analysis of DE genes in keratinocytes with miR-129 overexpression.

Table S4. DE genes in keratinocytes with circASH1L silencing.

Table S5. GO analysis of DE genes in keratinocytes with circASH1L silencing.

Table S6. Human sample information.

Table S7. List of reagents used in this study.

Table S8. List of growth factors and cytokines used in the study.

