## Supplemental Information for "Circular RNA circASH1L(4,5) protects microRNA-129-5p from target-directed microRNA degradation in human skin wound healing"

**This file includes the following:**

Figures S1 to S6

Tables S6 to S8

**Other supplemental materials not included in this file:**

Table S1 to S5

**Supplemental figures**

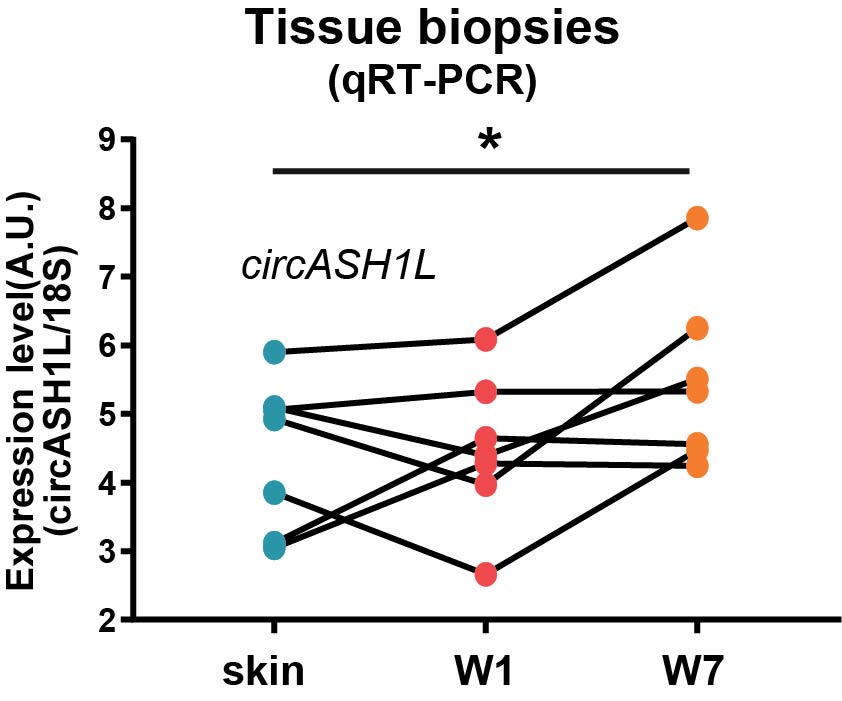

**Figure S1. qRT-PCR analysis of circASH1L in human skin, day 1 (W1), and day 7 wound (W7) tissues** (n = 7 donors). **P* < 0.05; One-way ANOVA and multiple comparisons test.

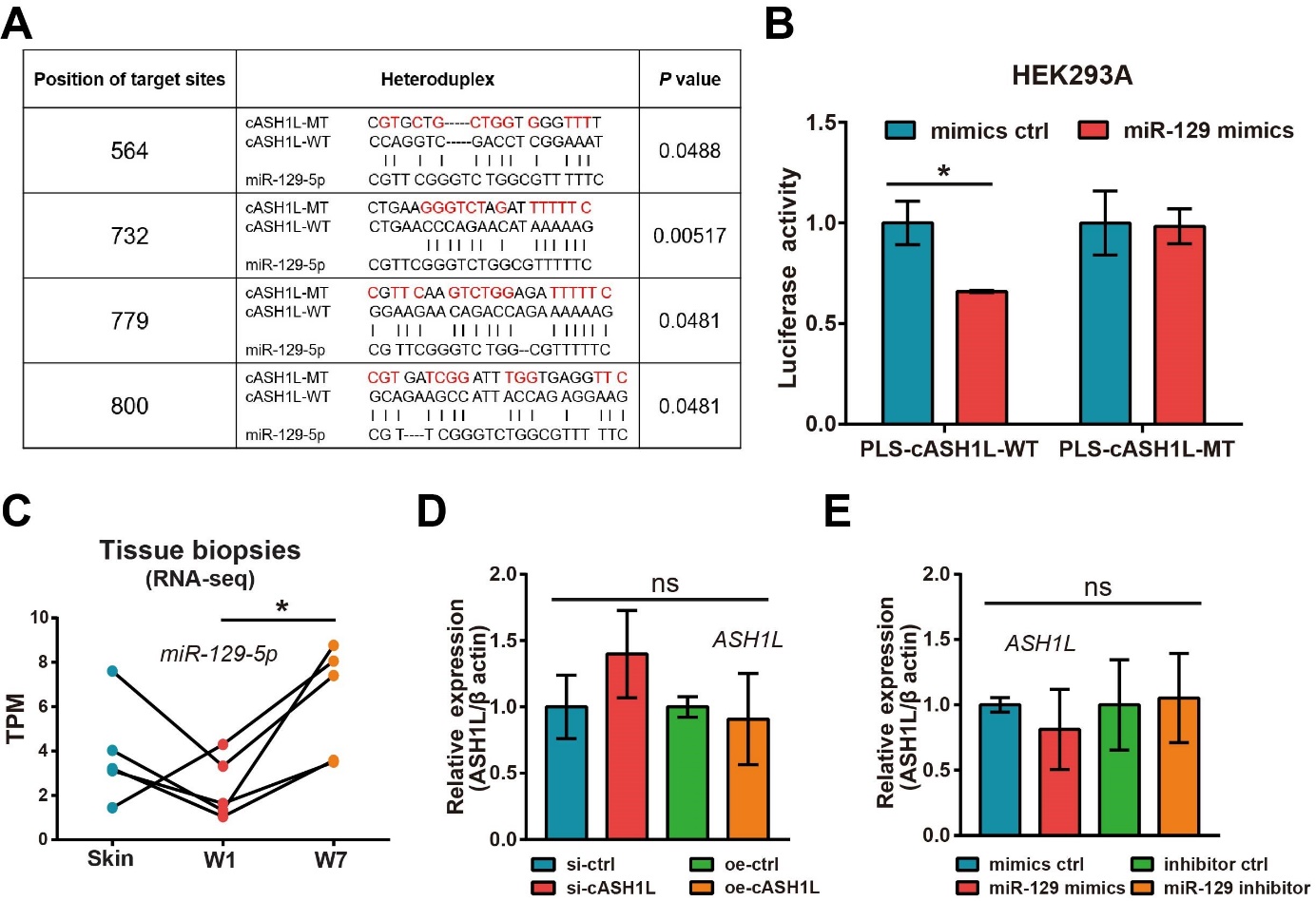
**Figure S2. CircASH1L binds to miR-129-5p and increases its level in human wound-edge keratinocytes.**

(A) Predicted miR-129-5p binding sites at circASH1L. The sequence of mutated miR-129 binding sites (MT) is highlighted in red.

(B) Luciferase assays were conducted using HEK293A cells transfected with luciferase reporter plasmids containing wild type (WT) or mutant type (MT) of circASH1L sequence together with miR-129 mimics or control oligos (n=4).

(C) RNA-seq analysis of miR-129-5p in human skin, day 1 (W1), and day 7 wound (W7) tissues (n=5).

(D and E) qRT-PCR analysis of ASH1L mRNA in HEKa cells transfected with circASH1L siRNAs (si-cASH1L) or overexpressing plasmids (oe-cASH1L) or their controls (si-ctrl or oe-ctrl) (n=4) (D); and in HEKa transfected with miR-129 mimics, inhibitors, or their controls (n=4) (E).

Data are presented as means ± SD or individual values; **P* < 0.05, ns: not significant; Unpaired Student’s t-test (B, D, E); One-way ANOVA and multiple comparisons test (C).

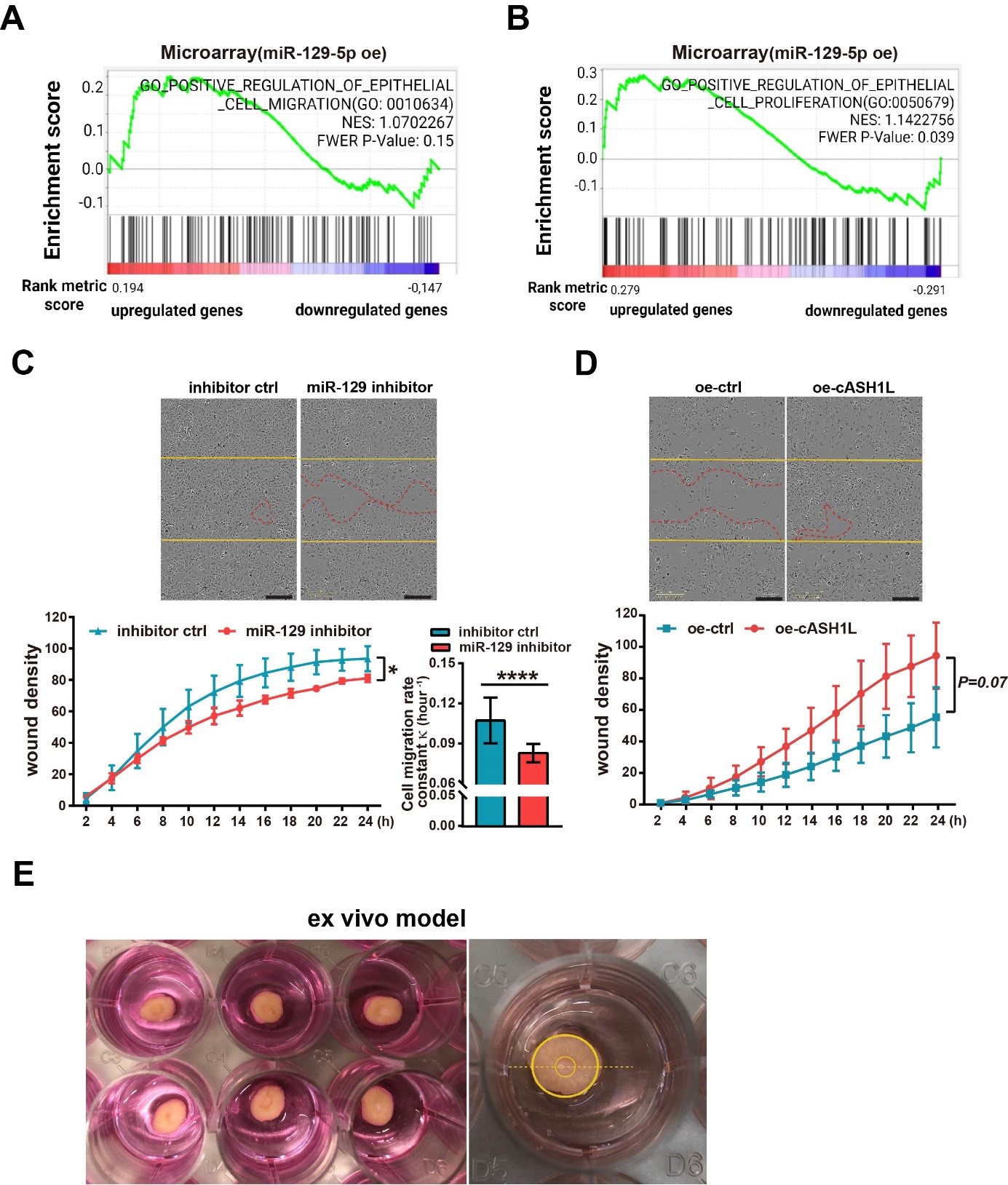

**Figure S3. CircASH1L-miR-129 axis promotes keratinocyte migration and proliferation.** (A and B) Gene set enrichment analysis (GSEA) of positive regulation of epithelial cell migration and proliferation in the microarray data of HEKa with miR-129-5p overexpression (oe).

(C and D) Scratch wound assays in HEKa cells with miR-129 inhibition (n=3) (C) or circASH1L overexpression (n=3) (D). Yellow and red lines indicate the wound-edges at 0 and 24hours, respectively. Scale bar = 300 μm.

(E) Human *ex vivo* wound model in culture: we sliced each biopsy at the wound's center, indicated by a yellow dotted line, before embedding them in paraffin for sectioning.

Data are presented as means ± SD; **P* < 0.05; *****P* < 0.0001; Two-way ANOVA and multiple comparisons test (C, D); Unpaired Student’s t-test (C).

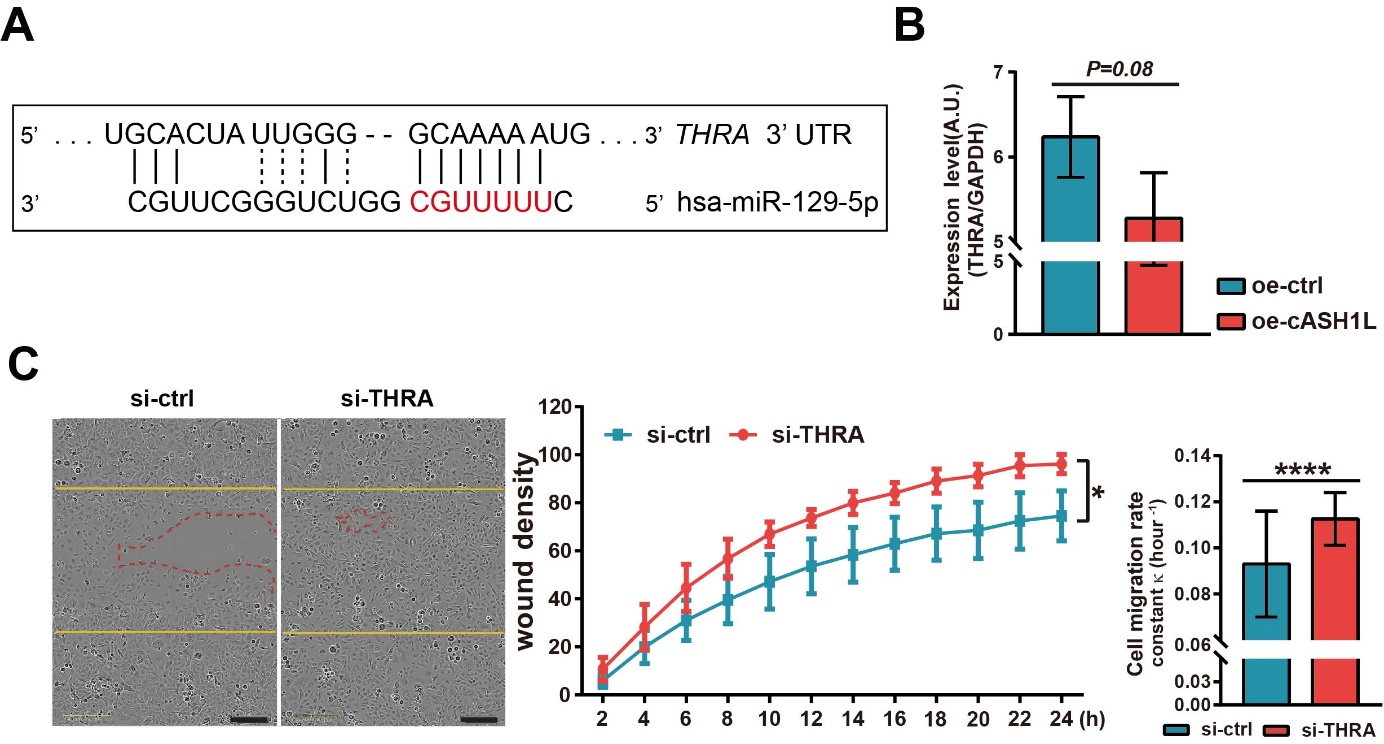
**Figure S4. miR-129-5p promotes keratinocyte migration by inhibiting THRA.**

(A) Illustration of miR-129-5p binding site at the 3'UTR of THRA.

(B) qRT-PCR analysis of THRA in HEKa cells with circASH1L overexpression (oe, n=3).

(C) Scratch wound assay in HEKa cells transfected with si-ctrl or si-THRA (n=4). Yellow and red lines indicate the wound-edges at 0 and 24hours, respectively. Scale bar = 300 μm.

Data are presented as means ± SD; **P* < 0.05; *****P* < 0.0001; Unpaired Student’s t-test (B, C); Two-way ANOVA and multiple comparisons test (C).

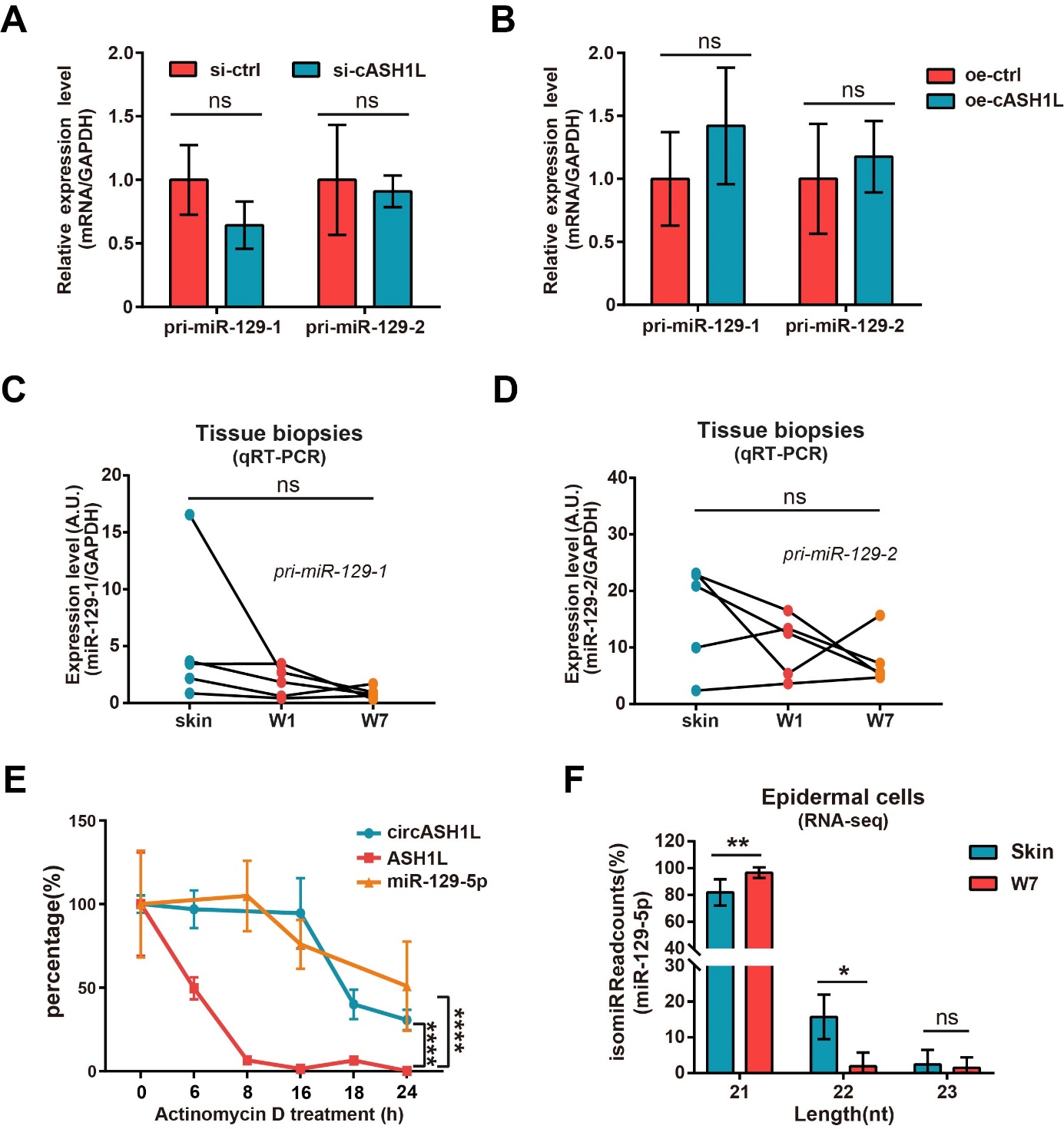

**Figure S5. CircASH1L enhances miR-129 stability.**

(A and B) qRT-PCR analysis of pri-miR129-1 and pri-miR129-2 in HEKa cells with circASH1L silencing (A) and overexpression (B) (n=4).

(C and D) qRT-PCR of pri-miR129-1 and pri-miR129-2 confirmed the expression level changes of pri-miR129-1, pri-miR129-2 in human skin, day 1 (W1), and day 7 wound (W7) tissues (n=5).

(E) qRT-PCR analysis of circASH1L, ASH1L mRNA, and miR-129-5p in HEKa cells treated with actinomycin D for 0-24hours (n=4).

(F) Analysis of the length of miR-129-5p isoforms in the small RNA-seq data of human skin and day-7 wound (W7) epidermal cells.

Data are presented as means ± SD or individual values; **P* < 0.05; ***P* < 0.01; *****P* < 0.0001; Two-way ANOVA and multiple comparisons test (A, B, E, F); One-way ANOVA and multiple comparisons test (C, D).

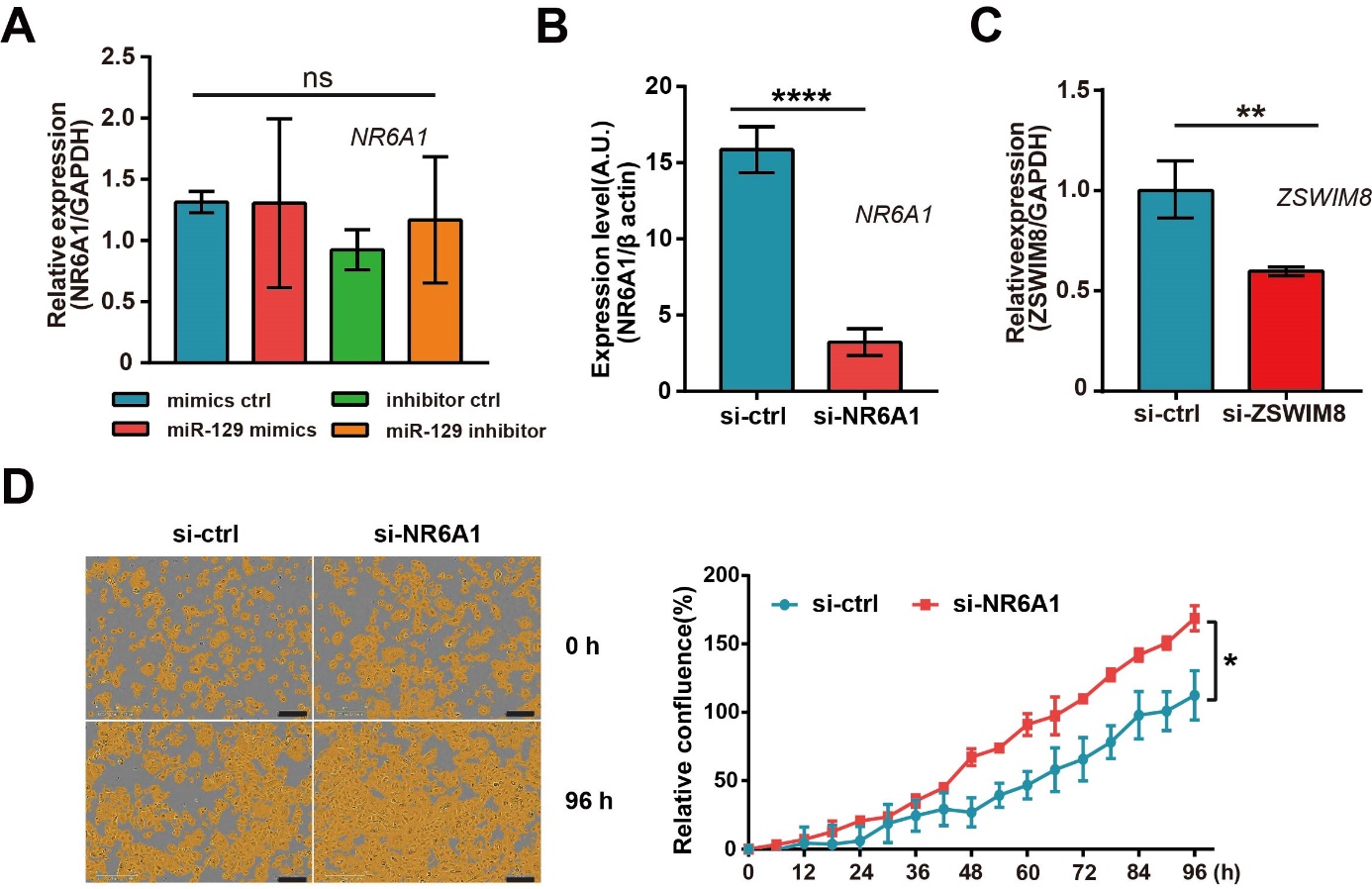
**Figure S6. NR6A1 is a TDMD trigger and suppresses keratinocyte growth.**

(A) qRT-PCR of NR6A1 in HEKa with miR-129-5p knockdown or overexpression (n=4).

(B) qRT-PCR of NR6A1 in HEKa transfected with NR6A1 siRNA or control siRNA (n=4).

(C) qRT-PCR of ZSWIM8 in HEKa transfected with ZSWIM8 siRNA or control siRNA (n=4). (D) Proliferation assays in HEKa cells transfected with si-NR6A1 (n=4). Scale bar = 300 μm. Data are presented as means ± SD; **P* < 0.05; ***P* < 0.01; *****P* < 0.0001; Unpaired Student’s t-test (A, B, C); Two-way ANOVA and multiple comparisons test (D).

**Supplemental tables**

**Table S6. Human sample information**

| **Donor** | **Sex** | **Age** | **Ethnicity** | **Body location** | **Experiments** |
| --- | --- | --- | --- | --- | --- |
| 1 | F | 66 | Caucasian | Lower leg | RNA-seq |
| 2 | M | 69 | Caucasian | Lower leg | RNA-seq |
| 3 | F | 67 | Caucasian | Lower leg | RNA-seq |
| 4 | M | 69 | Caucasian | Lower leg | RNA-seq, qRT-PCR |
| 5 | F | 64 | Caucasian | Lower leg | RNA-seq, qRT-PCR |
| 6 | F | 60 | Caucasian | Lower leg | qRT-PCR |
| 7 | F | 66 | Caucasian | Lower leg | qRT-PCR |
| 8 | F | 60 | Caucasian | Lower leg | qRT-PCR |
| 9 | F | 67 | Caucasian | Lower leg | qRT-PCR |
| 10 | F | 65 | Caucasian | Lower leg | qRT-PCR |
| 11 | M | 26 | Caucasian | Lower back | RNA-seq: keratinocyte |
| 12 | F | 30 | Caucasian | Lower back | RNA-seq: keratinocyte |
| 13 | M | 45 | Caucasian | Lower back | RNA-seq: keratinocyte |
| 14 | F | 43 | Caucasian | Lower back | RNA-seq: keratinocyte |
| 15 | M | 22 | Caucasian | Lower back | RNA-seq: keratinocyte |
| 16 | F | 24 | Caucasian | Breast | *Ex vivo* wound model |
| 17 | F | 43 | Caucasian | Abdominal | *Ex vivo* wound model |
| 18 | F | 33 | Caucasian | Abdominal | *Ex vivo* wound model |

F: Female, M: Male

**Table S7. List of reagents used in this study**

| **Primers/Probes** | **Sequence/Cat. no/Vendor** | |
| --- | --- | --- |
| CircASH1L(4,5) | F:GAAGACCTTTTCCGGGTAGG  R:GCAGAAGCCATTACCAGAGG | |
| hsa-miR-129-5p | TaqMan™ MicroRNA Assay ID 000590  (ThermoFisher Scientific) | |
| pri-hsa-miR-129-1 | TaqMan™ MicroRNA Assay ID Hs03302824_pri  (ThermoFisher Scientific) | |
| pri-hsa-miR-129-2 | TaqMan™ MicroRNA Assay ID Hs03303241_pri  (ThermoFisher Scientific) | |
| *RNU48* | TaqMan™ MicroRNA Assay ID 001006  (ThermoFisher Scientific) | |
| *ASH1L* | F: AAGAAGCCAGATGATGACACC  R: GTCTCCTATGTCTCCGTACTTA | |
| *NR6A1* | F: ATACACACATCAGCCGAACC  R: ACACTGGTCTTGCAGGAATG | |
| *THRA*  *GAPDH* | F: CCCTAGTTACCTGGACAAAGAC  R: ACGGCGATGCACTTCTT  F: GGTGTGAACCATGAGAAGTATGA  R: GAGTCCTTCCACGATACCAAAG  probe: AGATCATCAGCAATGCCTCCTGCA | |
| *β-actin*  *18S* | F: CATGTACGTTGCTATCCAGGC  R: CTCCTTAATGTCACGCACGAT  F: CGGCTACCACATCCAAGGAA  R: GCTGGAATTACCGCGGCT | |
| **siRNAs/mimics/inhibitor** | **Sequence/Cat. no/Vendor** | |
| si-circASH1L_01 | (AAG AGA AUA AUA AAA GGC UCC)TT | |
|  | siRNA fully targeting circASH1L BSJ; Dharmacon™ | |
| si-circASH1L_02 | (GAG AAU AAU AAA AGG CUC CCA)TT  siRNA fully targeting circASH1L BSJ; Dharmacon™ | |
| si-circASH1L_03 | (AGA GAA UAA UAA AAG GCU CCC)TT | |
|  | siRNA fully targeting circASH1L BSJ; Dharmacon™ | |
| si-ctrl | D-001210-03-05  siGENOME Non-Targeting siRNA; Dharmacon™ | |
| miR-129-5p mimics | C-300538-05-0005 miRIDIAN microRNA Human hsa-miR-129-5p - Mimic; Dharmacon™ | |
| mimics ctrl | CN-001000-01 miRIDIAN microRNA Mimic Negative Control; Dharmacon™ | |
| miR-129-5p inhibitor | IH-300539-05-0005 miRIDIAN microRNA Human  hsa-miR-129-5p Hairpin Inhibitor; Dharmacon™ | |
| inhibitor ctrl | IN-002005-01-05 miRIDIAN microRNA Hairpin Inhibitor Negative Control; Dharmacon™ | |
| si-THRA ON-TARGET™plus | L-003446-00-0005  siRNA targeting THRA; Dharmacon™ | |
| si-NR6A1 ON-TARGET™plus | L-003431-00-0005  siRNA targeting NR6A1; Dharmacon™ | |
| **Plasmids** | **Vendor/Information** | **Experiment** |
| circASH1L+up/down 200bp sequence | Eurofins (# 07074358) | circASH1L overexpression |
| Upstream intron(814bp) | Eurofins (# 07074358) |  |
| Downstream intron (813 bp) | Eurofins (# 07074358) |  |
| pcDNA3.1(+) (backbone) | Addgene (# V790-20) |  |
| pEX-A128-circASH1L_WT | Eurofins (# 06968576) | Luciferase reporter assay |
| pEX-A128-circASH1L_Mut | Eurofins (# 06945722) |  |
| 3’UTR Vector | Switchgear Genomics  (# S890005) |  |

**Table S8. List of growth factors and cytokines used in the study**

| **Growth factors**  **/Cytokines** | **Cat.no** | **Concentration** | **Vendor** |
| --- | --- | --- | --- |
| VEGF | 11343663 | 20ng/mL | ImmunoTools |
| FGF-2 | 11343625 | 30ng/mL | ImmunoTools |
| EGF | 11343406 | 20ng/mL | ImmunoTools |
| HBEGF | 259-HE-050 | 20ng/mL | ImmunoTools |
| KGF | 11343653 | 20ng/mL | R&D Systems |
| IGF | 11343314 | 20ng/mL | ImmunoTools |
| TGF-β1 | 11343161 | 10ng/mL | ImmunoTools |
| TGF-β2 | 11344751 | 10ng/mL | ImmunoTools |
| TGF-β3 | 11344482 | 10ng/mL | ImmunoTools |
| IL-1⍺ | 11349013 | 20ng/mL | ImmunoTools |
| IL-6 | 11340060 | 50ng/mL | ImmunoTools |
| IL-23A | 11340232 | 10ng/mL | ImmunoTools |
| IL-36⍺ | 11340362 | 10ng/mL | ImmunoTools |
| TNF⍺ | 210-TA | 50ng/mL | R&D Systems |
| MCP-1 | 11343380 | 10ng/mL | ImmunoTools |
| GM-CSF | 11343123 | 50ng/mL | ImmunoTools |
